# Unified establishment and epigenetic inheritance of DNA methylation through cooperative MET1 activity

**DOI:** 10.1101/2022.09.12.507517

**Authors:** Amy Briffa, Elizabeth Hollwey, Zaigham Shahzad, Jonathan D. Moore, David B. Lyons, Martin Howard, Daniel Zilberman

**Author notes:** These authors contributed equally to this work (Amy Briffa, Elizabeth Hollwey).

## Abstract

Methylation of CG dinucleotides (mCG), which regulates eukaryotic genome functions, is epigenetically propagated by Dnmt1/MET1 methyltransferases. How mCG is established and transmitted across generations despite imperfect enzyme fidelity remains mysterious. Here we show that MET1 *de novo* activity, which is enhanced by existing proximate methylation, seeds and stabilizes mCG in *Arabidopsis thaliana* genes. MET1 activity is restricted by active demethylation and suppressed by histone variant H2A.Z. Based on these observations, we develop a mathematical model that precisely recapitulates mCG inheritance dynamics and predicts intragenic mCG patterns and their population-scale variation given only CG site spacing as input. The model reveals that intragenic mCG undergoes large, millennia-long epigenetic fluctuations, and can therefore mediate evolution on this timescale. Our results demonstrate how genic methylation patterns are created, reconcile imperfect mCG maintenance with long-term stability, and establish a quantitative model that unifies the establishment and epigenetic inheritance of mCG.

**Highlights:**

- MET1 mediates a unified process of mCG establishment and maintenance within genes
- ROS1 and H2A.Z regulate the epigenetic dynamics of genic mCG
- A mathematical model predicts genic mCG patterns and their population variance
- Genic mCG undergoes large epigenetic fluctuations that can last thousands of years

## Introduction

Methylation of the fifth carbon of cytosine is the most common and extensively studied DNA modification in eukaryotes. In many lineages, including plants and animals, methylation is most prevalent within CG sites (Schmitz et al., 2019). CG methylation (mCG) can be epigenetically inherited over various timescales (Catania et al., 2020; Fitz-James & Cavalli, 2022; Petryk et al., 2021; van der Graaf et al., 2015; Vidalis et al., 2016), and its patterns are important for plant and animal development, and for human health (Greenberg & Bourc’his, 2019; Kumar & Mohapatra, 2021; Zhang et al., 2018). Multiple diseases, including cancer, are associated with mCG changes (Baylin & Jones, 2016; Jin & Liu, 2018). Mammalian age can be accurately predicted based on mCG patterns, indicating that the epigenetic inheritance and reprogramming of mCG is intimately associated with the aging process (Horvath & Raj, 2018). Understanding how mCG is epigenetically inherited, and how errors in this process are accommodated, is therefore crucial.

DNA methylation is classically divided into the establishment and maintenance phases (Chen & Riggs, 2011; Law & Jacobsen, 2010; Petryk et al., 2021). Establishment installs methylation at an unmethylated locus, and involves so-called *de novo* DNA methyltransferases, which belong to the Dnmt3 family in plants and animals (Law & Jacobsen, 2010). The establishment process is not itself epigenetic, but is determined, for example, by expression of a *de novo* DNA methyltransferase or an accessory factor at a specific time in a specific tissue (Bourc’his et al., 2001; Long et al., 2021). In flowering plants, the Dnmt3 (DRM) enzymes are guided by small RNA molecules within the RNA-directed DNA methylation (RdDM) pathway (Erdmann & Picard, 2020; Matzke & Mosher, 2014). RdDM establishes cytosine methylation in all sequence contexts, including CG, within transposable elements (TEs) and other repetitive sequences (Erdmann & Picard, 2020; Matzke & Mosher, 2014).

In the maintenance phase, the methylated state is epigenetically reproduced across cell divisions and – at least in plants – across multicellular generations (Niederhuth & Schmitz, 2014; Petryk et al., 2021). CG methylation is maintained by Dnmt1 enzymes in land plants and animals, and by Dnmt5 in several other eukaryotic lineages (Catania et al., 2020; Huff & Zilberman, 2014; Tirot et al., 2021). These enzymes use a semiconservative mechanism based on CG site symmetry. DNA replication produces a CG site methylated only on the template strand – a state that is recognized and converted to a fully methylated state by Dnmt1 or Dnmt5 (Dumesic et al., 2020; Hashimoto et al., 2008; Wang et al., 2022). This maintenance reaction has a similar error rate for Dnmt5 in the fungus *Cryptococcus neoformans* and for MET1 (the plant version of Dnmt1) in *Arabidopsis thaliana*, suggesting a fundamental limit on mCG maintenance fidelity (Becker et al., 2011; Catania et al., 2020; Schmitz et al., 2011; van der Graaf et al., 2015).

Mechanistic understanding of DNA methylation dynamics has been substantially aided by mathematical models (Busto-Moner et al., 2020; De Riso et al., 2020; Goyal et al., 2006; Haerter et al., 2014; Lövkvist et al., 2016; Rulands et al., 2018; Sontag et al., 2006; Zagkos et al., 2019). All existing mathematical models of mCG epigenetic inheritance employ *de novo* methylation – untemplated methylation of fully unmethylated CG sites – to compensate for semiconservative maintenance failure. These models therefore underline that the distinction between establishment and epigenetic maintenance of methylated regions is different to the one between *de novo* and semiconservative methylation at individual CG sites, because both of the latter are thought to be involved in the maintenance of a methylated region (Jeltsch & Jurkowska, 2014; Johannes & Schmitz, 2019). The models also emphasize strong cooperativity between nearby CG sites, such that methylation at one site can, for example, enhance methylation gain at nearby CG sites (Haerter et al., 2014; Lövkvist et al., 2016; Sontag et al., 2006; Wang et al., 2020). This feedback underpins bistability – the capacity for stable epigenetic inheritance of either the methylated or the unmethylated state at the same sequence under identical conditions (Haerter et al., 2014; Lövkvist et al., 2016; Zagkos et al., 2019).

In *Arabidopsis* TEs, MET1-catalyzed mCG maintenance is complemented by the RdDM pathway, as well as by the chromomethylase (CMT) pathway, which relies on a feedback loop between the CMT2 & 3 DNA methyltransferases and the SUVH4, 5 & 6 histone methyltransferases (Du et al., 2012, 2015; Rajakumara et al., 2011). The CMT pathway is attracted by mCG, and DNA methylation catalyzed by the CMT enzymes attracts RdDM, so that existing mCG promotes CMT and RdDM activity (To & Kakutani, 2022). The RdDM and CMT pathways together drive untemplated, *de novo* mCG at unmethylated sites at rates that far exceed the rate of MET1 maintenance failure, thereby stabilizing mCG inheritance (Lyons et al., 2022).

Flowering plants and most invertebrates have a type of intragenic methylation, called gene body methylation (gbM), that is distinct from TE methylation (Muyle et al., 2022). GbM tends to occur in exonic nucleosomes of conserved, constitutively expressed genes (Chodavarapu et al., 2010; Feng et al., 2010; Lewis et al., 2020; Zemach et al., 2010), and can prevent aberrant intragenic transcription and promote gene expression in *Arabidopsis* (Choi et al., 2020; Shahzad et al., 2021). Compared to TE methylation, the epigenetics of gbM are far more mysterious (Muyle et al., 2022). GbM is characterized by the absence of non-CG methylation (the type catalyzed by the DRM and CMT enzymes) (Takuno & Gaut, 2012; Zemach & Zilberman, 2010) and by a more equal balance between the rates of mCG loss and gain (van der Graaf et al., 2015). The RdDM pathway is thought not to play a major role in gbM establishment (Vaughn et al., 2007). Instead, CMT3 has been proposed as the main establishment enzyme, but the only direct evidence for this is from experiments in which CMT3 is ectopically overexpressed (Bewick et al., 2016; Wendte et al., 2019). Hence, the gbM establishment mechanism and the source of *de novo* mCG during epigenetic maintenance remain unknown. How apparently random mCG losses and gains at individual CG sites nonetheless produce gbM patterns that remain coherent over long periods of time is also unknown. Why gbM favors nucleosomes and exons is unclear, and why some genes are reproducibly methylated across time, others are reproducibly unmethylated, and yet others are variably methylated is mysterious (Kawakatsu et al., 2016; Niederhuth et al., 2016; Zhang et al., 2020).

Here, we comprehensively elucidate gbM epigenetics using *Arabidopsis* as a model. We show that gbM establishment, maintenance, and even loss, constitute a unified process mediated by MET1. MET1 activity is suppressed by the histone variant H2A.Z and, especially in unmethylated genes, by the DNA demethylase ROS1. MET1 *de novo* and maintenance activities are enhanced by proximate mCG and are therefore not spatially homogeneous. We use a mathematical model to precisely reproduce the *in vivo* patterns of mCG loss and gain observed over experimentally assessable timescales (up to 30 generations). The same model, given only CG site spacing as input, accurately predicts steady state gbM patterns – including the preference for exons and nucleosomes – and their variance across *Arabidopsis* natural populations. The steady-state prediction is independent of the initial methylation state, implying a lack of bistability, whereas any level of methylation is stably inherited over short timescales. Steady state gbM is subject to stochastic epigenetic fluctuations that can be large and can last for thousands of years, a feature that establishes gbM as an epigenetic genotype able to mediate evolution on this timescale. Overall, we present a precise, quantitative and predictive understanding of gbM epigenetics that far surpasses that of any other epigenetic process.

## Results

### GbM epigenetic dynamics are primarily mediated by MET1

To investigate the epigenetic inheritance of mCG within genes, we grew *Arabidopsis* for up to six consecutive generations (Fig. S1A) and obtained whole-genome bisulfite sequencing data. In addition to wild type (*WT*) Col-0, we analyzed mutants of all non-MET1 methyltransferases in different combinations (*cmt2, cmt3, drm1drm2, cmt2cmt3, drm1drm2cmt2cmt3* (*ddcc*)). We identified single CG site methylation gains and losses at each generation and calculated epimutation rates per cell cycle based on the published estimate of 34 cell cycles per generation (Fig. 1A-C; Fig. S1B; Table S1-2) (Watson et al., 2016). Our rates for all *Arabidopsis* genes agree with those previously published (Fig. 1A; Table S1) (van der Graaf et al., 2015). Although a recent study concluded that sparsely methylated regions (SPMRs) within gbM genes have an enhanced epimutation rate (Hazarika et al., 2022), we find that epimutation rates are similar between SPMRs and methylated regions of gbM genes that lie outside SPMRs (Fig. S2A-D; Table S1-2).

**Figure 1.**
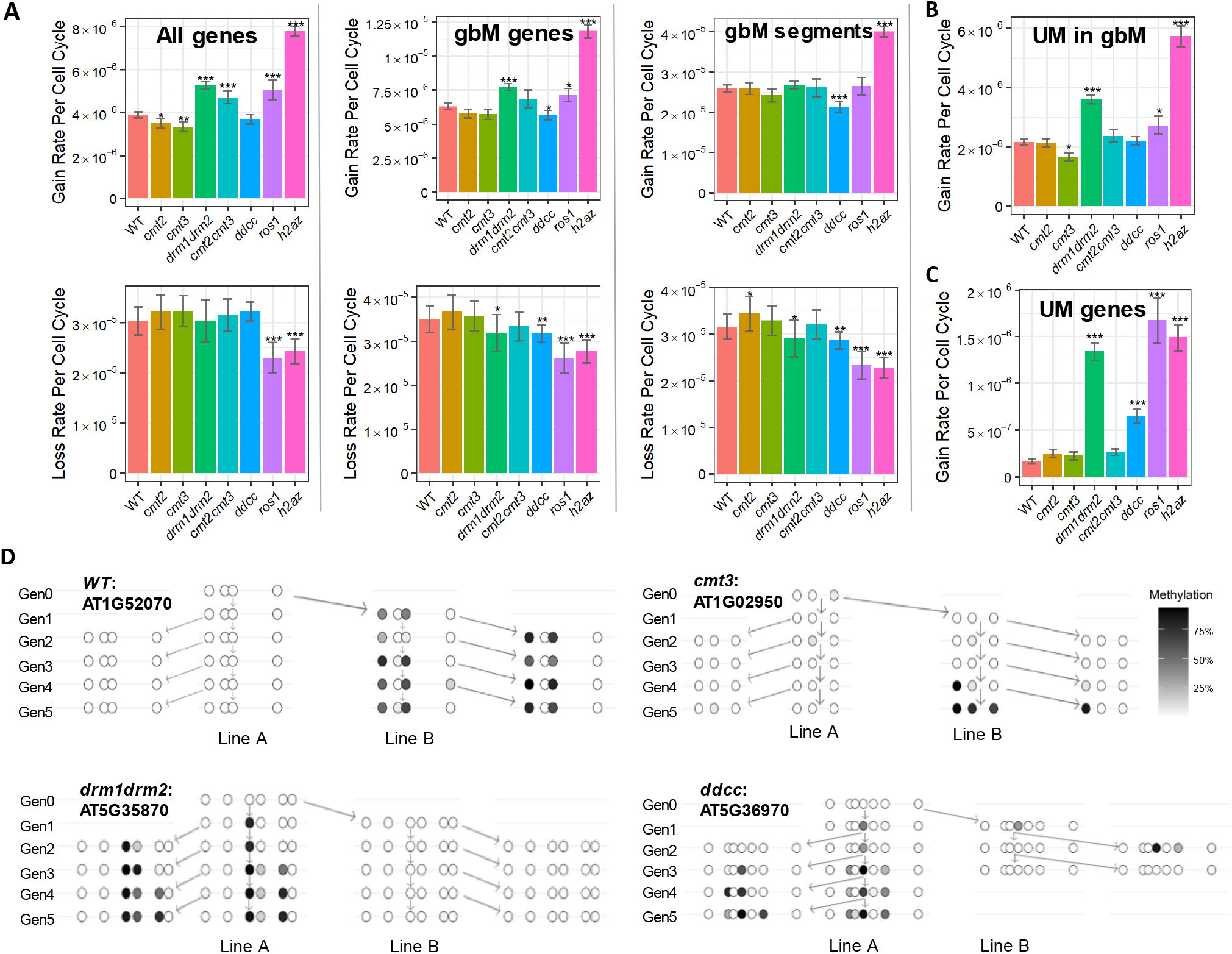
MET1 is a *de novo* methyltransferase. **A-C)** Per cell cycle rates of mCG gain and loss at individual CG sites within indicated genomic regions. A significant difference from the *WT* rate is indicated by *p=0.01 to 0.001, **p=0.001 to 0.00001, and ***p<0.00001 (Fisher’s exact test). Error bars indicate standard error. **D)** GbM gains that expand to form new methylation clusters can be observed in previously entirely unmethylated genes in different genotypes, including *ddcc*. Each circle represents an individual CG pair, with darkness of fill indicating fractional mCG.

We find that rates of mCG gain and loss are similar to *WT* in all the methyltransferase mutants we tested, including the quadruple *ddcc* mutant where the only functional methyltransferase is MET1 (Fig 1A-B; Table S1) (He et al., 2022). Reanalysis of published *cmt3* and *suvh4/5/6* triple mutant data (Hazarika et al., 2022) produced similar results (Fig. S2E-F). Nevertheless, rates are somewhat changed in some mutants, most significantly a decreased gain rate in gbM segments of *ddcc* plants and elevated gain rates in all genes, gbM genes and in unmethylated (UM) segments of gbM genes in *drm1drm2* mutants (Fig. 1A-B; Table S1). These differences likely reflect small contributions of non-MET1 methyltransferases to gbM, which may contribute to gbM steady state over long periods of time. However, our data demonstrate that gbM gains and losses are primarily due to MET1 activity (Fig 1A-B; Table S1), indicating that MET1 is both a maintenance and a *de novo* methyltransferase.

The *WT* mCG loss rate in genes is 25-fold higher than in TEs (Fig. 1A; Fig. S1B; Table S1), which is consistent with published results (van der Graaf et al., 2015). We observe significantly elevated TE loss rates in all methyltransferase mutants, and loss rates in *ddcc* mutants (2.0 × 10^-5^ per site per cell cycle) that are 18-fold higher than *WT* and similar to those in *WT* gbM segments (3.2 × 10^-5^ per site per cell cycle; Fig. 1A; Fig. S1B; Table S1). MET1 maintenance efficiency therefore appears to be comparable between genes and TEs in the absence of RdDM and CMT activity. TE mCG gain rates cannot be accurately measured using our (or analogous published) data because the measurements rely on ancestral UM sites that are rare and unrepresentative in TEs (Lyons et al., 2022). However, it is notable that we find TE mCG gains in all methyltransferase mutant lines, including *ddcc* (Fig. S1B), indicating that MET1 functions as a *de novo* methyltransferase in TEs as well as in genes.

### MET1 is a *bona fide de novo* methyltransferase

Methylation dynamics within gbM genes (and methylated TEs) may differ from those of unmethylated loci. To explore whether MET1 can establish methylation in such loci, we calculated mCG gain rates in unmethylated (UM) genes (Fig. 1C; Table S1). We observed no reduction of the gain rate in any of the analyzed mutants, and identified new methylation events in all methyltransferase mutants, including the quadruple *ddcc* mutant (Fig. 1C; Table S1). All genotypes except *cmt2*, including *ddcc*, also contain at least one example of a single mCG gain within a UM gene expanding to produce a new mCG cluster (Fig. 1D; Fig. S3; Table S3). These results demonstrate that methyltransferases other than MET1 are not required to initiate or expand gbM clusters. MET1 is therefore a true *de novo* methyltransferase that can stochastically establish gbM in previously unmethylated genes.

GbM patterns vary substantially between natural *Arabidopsis* accessions (Dubin et al., 2015; Kawakatsu et al., 2016; Pignatta et al., 2014; Schmitz et al., 2013). These differences could arise, at least in part, due to the stochastic *de novo* activity of MET1. To investigate this hypothesis, we determined whether UM genes that gained methylation in our experiments (all lines belong to the Col-0 accession), or in analogous published experiments, are more likely to be methylated in the *Arabidopsis* population than all Col-0 UM genes. Col-0 UM genes that gained gbM in our *WT* data have gbM in 46% of accessions on average (Table S4). The corresponding percentages for other genotypes are 50% for *cmt2*, 45% for *cmt3*, 42% for *cmt2cmt3*, 41% for published *WT* data (MAL), 32% for *drm1drm2*, and 35% for *ddcc* (Table S4). In comparison, all Col-0 UM genes are much less likely to contain gbM in the population (16%, p < 2×10^-16^, Table S4-5). This indicates that some UM Col-0 genes are predisposed to gain gbM due to *de novo* MET1 activity, and suggests that natural *Arabidopsis* gbM diversity reflects an accumulated pattern of stochastic differences that are initiated and epigenetically maintained by MET1.

### ROS1 preferentially prevents methylation gain in unmethylated gene bodies

The above results raise the question of why some genes are more likely to experience *de novo* MET1 activity. Two of the methyltransferase mutants we analyzed, *drm1drm2* and *ddcc*, exhibit significantly increased rates of mCG gain in UM genes (Fig. 1C; Table S1). We hypothesized that this may be due to reduced expression of the DNA demethylase *ROS1* that occurs in mutants of the RdDM pathway (Huettel et al., 2006; Penterman et al., 2007). We therefore analyzed mCG in *ros1* mutants over consecutive generations. We did not observe a significant change in the gain rate in gbM segments (Fig. 1A). The rate of mCG gain is elevated by about 25% in UM segments of gbM genes, and increases 10-fold in UM genes, to about the *WT* level in UM segments of gbM genes (Fig. 1B-C; Table S1). The loss rate in gbM segments is also reduced by about 25% in *ros1* mutants (Fig. 1A; Table S1). This indicates that ROS1 prevents unmethylated genes from gaining methylation and affects the mCG loss and gain rates in gbM genes.

New gbM clusters arose in UM genes of *ros1* mutants, but unlike in the other genotypes, these are not restricted to genes with gbM in other accessions (Table S3). A new cluster arose in a gene (AT4G08865) that contains gbM in only 0.4% of accessions (Fig. 2A), and clusters arose in AT4G34419 (gbM in 3.4% of accessions) and AT5G63715 (gbM in 2.6% of accessions; Table S3). In contrast, no gene with a population gbM frequency below 13.4% gained a gbM cluster in other genotypes (Table S3), suggesting that ROS1 maintains a subset of genes in a perpetually unmethylated state. Indeed, mean population gbM frequencies in Col-0 UM genes that gain gbM in *ros1* (33%) and low-ROS1 genotypes (32% in *drm1drm2* and 35% in *ddcc*) are significantly lower than such frequencies in other lines (41% to 50%, p < 7.46×10^-09^; Table S4-5). Therefore, ROS1 regulates the relative probability of gbM gain, so that genes that are unlikely to gain gbM become more susceptible without ROS1 activity.

**Figure 2.**
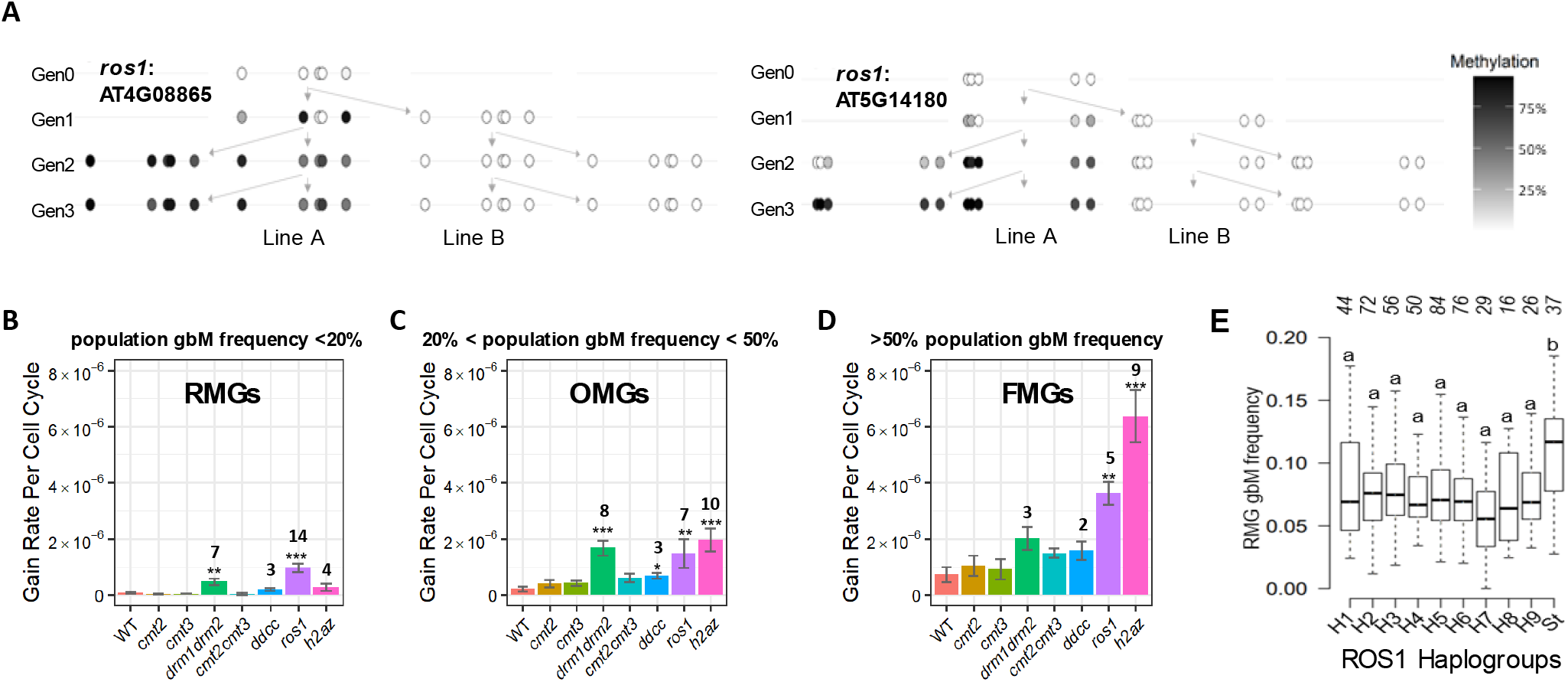
ROS1 and H2A.Z prevent mCG gain in unmethylated genes. **A)** New methylated clusters occur in entirely unmethylated genes in *ros1* mutants, including in a gene (AT4G08865) that is almost never methylated in the population. Each circle represents an individual CG pair, with darkness of fill indicating fractional mCG. **B-D)** Per cell cycle rates of mCG gain at individual CG sites in genes that are unmethylated (UM) in Col-0, grouped depending on their population gbM frequency into rarely methylated genes (RMGs) (**B**, <20% population gbM frequency), occasionally methylated genes (OMGs) (**C**, 20-50% population gbM frequency), and frequently methylated genes (FMGs) (**D**, >50% population gbM frequency). A significant difference from the WT rate is indicated by *p=0.01 to 0.001, **p=0.001 to 0.00001, and ***p<0.00001 (Fisher’s exact test). Numbers above bars indicate fold-change from *WT* for genotypes that are significantly changed in UM genes overall. Error bars indicate standard error. **E)** Frequency of gbM within RMGs in ROS1 haplogroups based on the amino acid sequence, and an additional ROS1-St group that contains accessions with a premature stop codon. Different letters indicate significant differences at p < 0.05 (one-way ANOVA), i.e. groups denoted with “a” are statistically different from “b”. The number of accessions is indicated for each haplogroup (top).

To further investigate the connection between ROS1 and likelihood of gbM gain, we subdivided Col-0 UM genes based on their population gbM frequency into rarely methylated (RMGs; gbM in <20% of accessions), occasionally methylated (OMGs; gbM in >20% and <50% of accessions) and frequently methylated genes (FMGs; gbM in >50% of accessions). *WT* and *cmt* mutant lines (*cmt2, cmt3, cmt2cmt3*) show a clear progression of mCG gain rates, which increase with population gbM frequency (Fig. 2B-D; Table S1). In contrast, gain rates in *ros1* mutants are more uniform, so that the relative gain rate increase between *WT* and *ros1* is greatest in RMGs (14-fold) compared to FMGs (5-fold) (Fig. 2B-D, Table S1). This supports the conclusion that the relative probability of gbM gain is dependent on ROS1.

The above results indicate that natural accessions with reduced ROS1 activity should have an overabundance of gbM in genes that are rarely methylated in the population. To test this hypothesis, we examined the association between ROS1 amino acid polymorphism and RMG gbM frequency across accessions. We defined nine ROS1 haplotypes, and a tenth group containing alleles with premature stop codons (Fig. S4A; Table S6). Accessions carrying a premature stop codon (ROS1-St) display a high frequency of RMG gbM (Fig. 2E). Furthermore, mixed model genome-wide association (GWA) analysis using the number of methylated RMGs as the phenotype identified several SNPs around ROS1 that are marginally associated with RMG gbM frequency (Fig. S4B). This analysis also detected a significant association (SNP Chr5:23536319; Fig. S4C) near ROS3, an RNA-binding protein that functions in the ROS1 pathway to mediate active DNA demethylation (Zheng et al., 2008). Based on amino acid sequence variation, we defined nine ROS3 haplogroups (H1-H9). GWA SNP Chr5:23536319 is linked with ROS3-H8, which is associated with high RMG gbM frequency (Fig. S4D-E; Table S6). Taken together, these results indicate that the ROS1 pathway protects UM genes from methylation, affects mCG dynamics in gbM genes, and that differential ROS1 activity determines the population frequency of gbM within each gene.

### Local cooperativity shapes MET1 *de novo* activity

We observed that gains of single mCG sites in unmethylated genes were followed either by reversion to an unmethylated state or by rapid expansion of a gbM cluster (Fig. 1D; Fig. 2A; Fig. S3). This suggests cooperative dynamics, where the rates of methylation change are influenced by nearby mCG sites (Fig. 3A) (De Riso et al., 2020; Haerter et al., 2014; Lövkvist et al., 2016; Sontag et al., 2006). To examine this behavior, we plotted the likelihood of methylation loss and gain within gbM genes relative to the nearest mCG site (Fig. 3B-C). In addition to our newly generated data, we analyzed data from previous studies that assessed the inheritance of DNA methylation in *WT Arabidopsis* mutation accumulation lines (MAL) over 30 generations (Table S1-2) (Becker et al., 2011; Schmitz et al., 2011). The published data produced the same general results as our *WT* data (Fig. 3B-C). Individual lines within the MAL datasets produced similar patterns, as did the two different MAL datasets overall (Fig. S5A-C).

**Figure 3.**
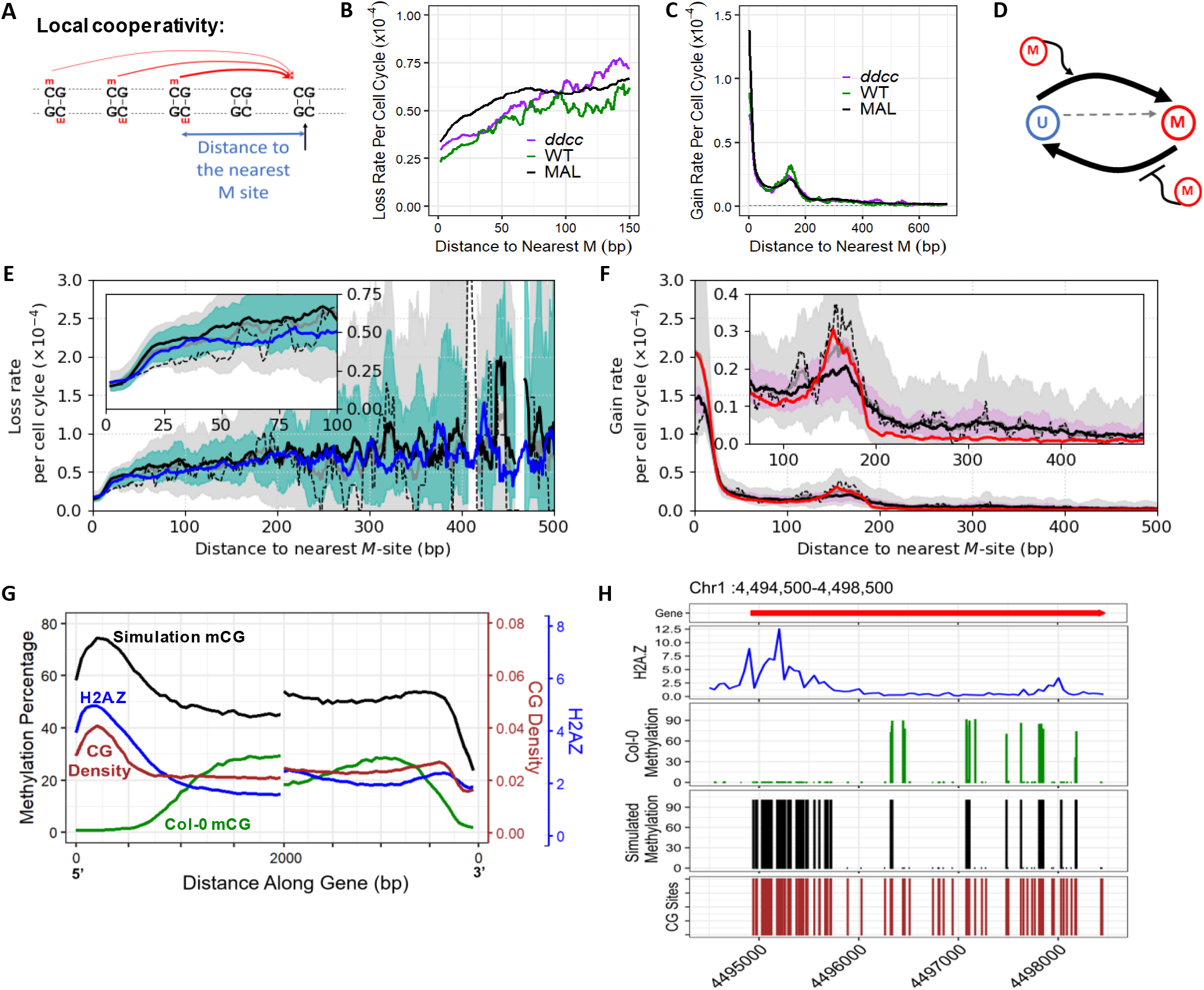
Existing methylation shapes mCG gain and loss. **A)** Schematic showing expected dependence of mCG gain rate on proximity to nearby methylated sites produced by a local cooperative interaction. Strength of the effect decreases with the distance between a mCG site and a target CG site. **B-C)** Data profiles of methylation loss (**B**) and gain (**C**) rates per cell cycle, plotted as a function of distance to the nearest mCG site, plotted over whole gbM genes. Gain rate plateaus at a level (5×10^-7^ per site per cell cycle; dashed line) comparable to the gain rate in entirely unmethylated genes. Rates shown as a 30 bp moving average. **D)** Schematic of the simulated effective two state model. **E-F)** Simulated profiles of mCG loss (**E**, blue line) and gain (**F**, red line) rates per cell cycle, plotted as a function of distance to the nearest mCG site, calculated over whole gbM genes, where each line is the mean of four simulated replicates (simulation time: 30 generations, initial state: experimental Col-0). Solid black lines represent the loss (**E**) and gain (**F**) rate profiles averaged over published MAL data (Schmitz et al., 2011), with the light-blue (**E**) and pink (**F**) bands corresponding to ±1 standard deviation (sd). Dotted black lines reproduce the Col-0 gain/loss rate profiles shown in **B** and **C**. Solid grey lines are mean gain/loss rates, averaged over all *WT* experimental data sets, with grey bands showing ±1 sd. Rates are shown as a 10 bp moving average. Model parameters given in Table S7A. Insets highlight loss rate at short length-scales (**E**) and enhanced gain rate at length-scale of ∼170bp (**F**). **G)** Methylation levels averaged over all gbM genes (N=14,581), with Col-0 mCG (green) and simulated mCG (a single modelling realization run for 100,000 generations from the Col-0 initial state; model parameters in Table S7A; black) (left y-axis). Mean CG density in brown and mean H2A.Z enrichment in blue (right y-axes). Left panel displays the 5’ end of the gene; right panel displays the 3’ end. **H)** Genome browser view of AT1G13180 with gene annotation in red. Col-0 mCG (green), simulated mCG (simulation as in **G**; black), Col-0 H2A.Z enrichment (blue) and positions of individual CGs to demonstrate CG density (red) are indicated.

Rates of gain and loss are shaped by proximity to mCG sites (Fig. 3B-C), with loss rates rising with increasing distance to other mCG sites (Fig. 3B). The effect of nearby mCG on methylation gain is even clearer, with the likelihood of gain over 100-fold above background within 25 bp of a methylated site, before dropping sharply (Fig. 3C). The methylation gain rate then rises again, peaking between 160-170 bp from the nearest mCG site, before plateauing to a background level (5×10^-7^ per site per cell cycle; indicated by the dotted line in Fig. 3C) by ∼1000 bp away from the nearest mCG site. The profiles of gain and loss rates are very similar between *WT* and all methyltransferase mutants, including the quadruple *ddcc* mutant (Fig. 3B-C; Fig. S5D). These results indicate that nearby mCG strongly stimulates MET1 *de novo* activity as well as either promoting MET1 maintenance activity or inhibiting DNA demethylation (or both).

### Cooperative gain and loss processes are required to model methylation dynamics

To gain a quantitative understanding of gbM dynamics, we developed a computational stochastic model (Fig. 3D; Fig. S6A). Because the experimental methylation gains around a given mCG site fall away as an approximate power law, the cooperative gain interaction strength for each uCG site is calculated using a power-law decay as a function of distance to existing mCG sites (Fig. S6B-C, Methods). Additionally, we include a uniform spontaneous gain rate for every uCG site. All CG sites within each gene are then initialized to an experimental Col-0 methylation state and simulated for 30 generations (1020 cell cycles) to match the published MAL data (Becker et al., 2011; Schmitz et al., 2011).

Although we could reproduce the rapid fall in gain rate at a length-scale of ∼30 bp, this simplest model fails to generate the second peak at ∼170 bp (Fig. S6D), demonstrating that this enhancement of the gain rate does not emerge from the intrinsic spatial distribution of CG sites. As ∼170 bp corresponds to the nucleosome repeat length (Zhang et al., 2015), we hypothesize that the 3D chromatin conformation reduces the effective distance between the target uCG and promoting mCG sites, thus generating an enhanced cooperative interaction when their separation matches the nucleosome repeat length. This motivated including a secondary, weaker, component to the cooperative gain interaction, with the origin of the peak centered on ∼170 bp (Fig. S6B).

The model also includes mCG loss, which may occur either through active demethylation (e.g., via ROS1) or passively due to maintenance failure at DNA replication. We find that a cooperative maintenance mechanism (by which surrounding mCG sites reduce the probability of a maintenance failure at the target mCG site) must be included to qualitatively reproduce the loss rate profile (Fig. S6C,E-F). Thus, our model contains four gain/loss processes: spontaneous *de novo*, cooperative *de novo*, background loss, and cooperative suppression of loss. This model quantitively recapitulates the experimental gain and loss profiles (Fig. 3E-F; Fig. S6G, Table S7A), indicating that a simple mathematical model with cooperative gains and losses can encapsulate complex epimutation patterns.

### Histone variant H2A.Z broadly reduces the rate of gbM gains

Because of the low rates of mCG loss and gain, a simulation over 30 generations only slightly alters the initial methylation pattern. To test whether the modelled epigenetic dynamics can reproduce the observed steady-state gbM patterns, we simulated entire genes for 100,000 generations (sufficient to reach steady-state) starting from the Col-0 methylation state, and calculated the average mCG level across each gene (Fig. S7A-B). Simulating entire gbM genes results in drastic over-methylation, as does simulating unmethylated genes. Conversely, simulating only Col-0 gbM segments produces under-methylation (Fig. S7C). The modelled steady-state methylation levels depend not only on the model parameters but also on the CG site number and density of the simulated regions. As a result, when simulating entire gbM genes, the model reproduces the observed decline of gbM toward the 3’ ends of genes (although the absolute level is too high). It fails, however, to reproduce the 5’ end decline, instead predicting a peak of mCG aligned with a corresponding enrichment of the CG dinucleotide density (Fig. 3G-H). Due to the nonlinear cooperative interaction, the over-methylation of 5’ gene ends drives higher average methylation of the entire gene (Fig. 3G-H). These findings indicate that only certain regions of some genes are subject to methylation, and that the epigenetic dynamics we measure and model only apply to these methylatable regions. This is consistent with our finding that ROS1 helps to protect constitutively unmethylated genes from methylation (Fig. 1C; Fig. 2A-D), and raises the question of what other biological processes might regulate methylation targeting within genes.

The regions that become over-methylated in our simulation (unmethylated genes and 5’ gene ends) are enriched for histone variant H2A.Z (Fig. 3G-H; Fig. S8A) (Zilberman et al., 2008). DNA methylation and H2A.Z are anticorrelated in plants and animals, and they can affect each other’s distribution (Fig. 3G-H) (Coleman-Derr & Zilberman, 2012; Conerly et al., 2010; Edwards et al., 2010; Murphy et al., 2018; To et al., 2020; Zemach et al., 2010; Zilberman et al., 2008). Consistently, the genes over-methylated by the model have relatively high H2A.Z, whereas the genes under-methylated by the model have relatively low H2A.Z compared to the accurately modelled genes (Fig. S8B). This suggests that gbM epigenetic dynamics are influenced by H2A.Z.

To investigate how H2A.Z shapes gbM, triple *hta8 hta9 hta11* (*h2az*) mutants were grown and analyzed over multiple generations. We observed a significant increase in the methylation gain rate in *h2az* compared to *WT*, with the strongest relative effect in UM genes (8.8-fold increase), followed by UM regions of gbM genes (2.6-fold increase) and gbM segments (1.5-fold increase; Fig. 1A-C; Table S1). Loss rates in gbM genes and segments decreased to about the same extent as in *ros1* mutants (Fig. 1A; Table S1). Thus, H2A.Z, like ROS1, preferentially lowers mCG gains in unmethylated sequences and increases mCG losses in gbM segments. However, unlike ROS1, H2A.Z has a significant effect on mCG gain in gbM segments (Fig. 1A; Table S1). Also, unlike in *ros1* mutants, Col-0 UM genes with methylation gains in *h2az* mutants have an average population gbM frequency (47%) similar to that in *WT* data (Table S4-5). Consistently, lack of H2A.Z causes a smaller (and non-significant) relative mCG gain rate increase in RMGs (4-fold) than in UM genes with higher population gbM frequencies (∼9-fold; Fig. 2B-D; Table S1). Therefore, H2A.Z does not preferentially influence RMGs as ROS1 does, but instead broadly suppresses gbM gain.

The findings that H2A.Z and ROS1 preferentially suppress mCG gains in UM sequences indicate that the chromatin environment and active DNA demethylation specify methylatable sequences within the genome. We therefore constructed an annotation of methylatable gene-regions. To account for the intrinsically stochastic methylation dynamics, which will at times create unmethylated regions even in locations that can in principle be methylated, we selected accessions with overall gbM levels similar to Col-0 (N=891) (Kawakatsu et al., 2016) and used these to define genes that could contain gbM. Accessions were considered similar to Col-0 if they have a global gbM level within one standard deviation of the mean, because independent Col-0 samples vary substantially within this interval (Fig. S9A). Gene ends were removed using the location of methylation in these accessions, and the level of H2A.Z in Col-0. This procedure produced a single, continuous, methylatable region per gene in 7980 genes (Table S8).

### The model can accurately fit steady-state gbM within methylatable regions

We updated model parameters to fit the gain/loss rates of methylatable regions (Table S7B) and tested whether the revised epigenetic dynamics can reproduce the observed steady-state gbM patterns. Average mCG levels over methylatable regions of 30 high-coverage Col-like accessions corresponded well with the steady-state mCG patterns produced by our model (100,000 generation simulations repeated 30 times to match the number of accessions; Fig. S9B). However, splitting the mCG level distributions by CG site density (Fig. S9C) revealed that the model systematically over-methylated CG-rich methylatable regions and under-methylated CG-poor regions.

To correct for this issue, we introduced a CG density dependent linear correction to the cooperative gain and cooperative maintenance strengths, such that higher densities led to reduced cooperativity per site. It is likely that the requirement for this correction is (at least in part) due to assuming that each mCG site independently enhances the cooperativity strength. Actually, however, the contribution of each individual site may be reduced when there are many mCG sites in close vicinity. With re-optimized parameters (Table S7C), we successfully achieved a quantitative fit simultaneously to the Col-0 30-generation gains/losses (Fig. S10A-C) and the steady-state average methylation level for methylatable regions of 30 Col-like accessions (Fig. 4A; Fig. S9C). Note that cooperative suppression of losses accounts for a ∼4-fold loss reduction over 30 generations, whereas cooperative promotion of gains accounts for ∼6-fold increase in gains (Table S9). This illustrates that although gains cooperativity is much stronger over short distances (Fig. 3B-C), cooperative gain and loss dynamics make comparable contributions to gbM epigenetic inheritance and are therefore of roughly equal importance.

**Figure 4.**
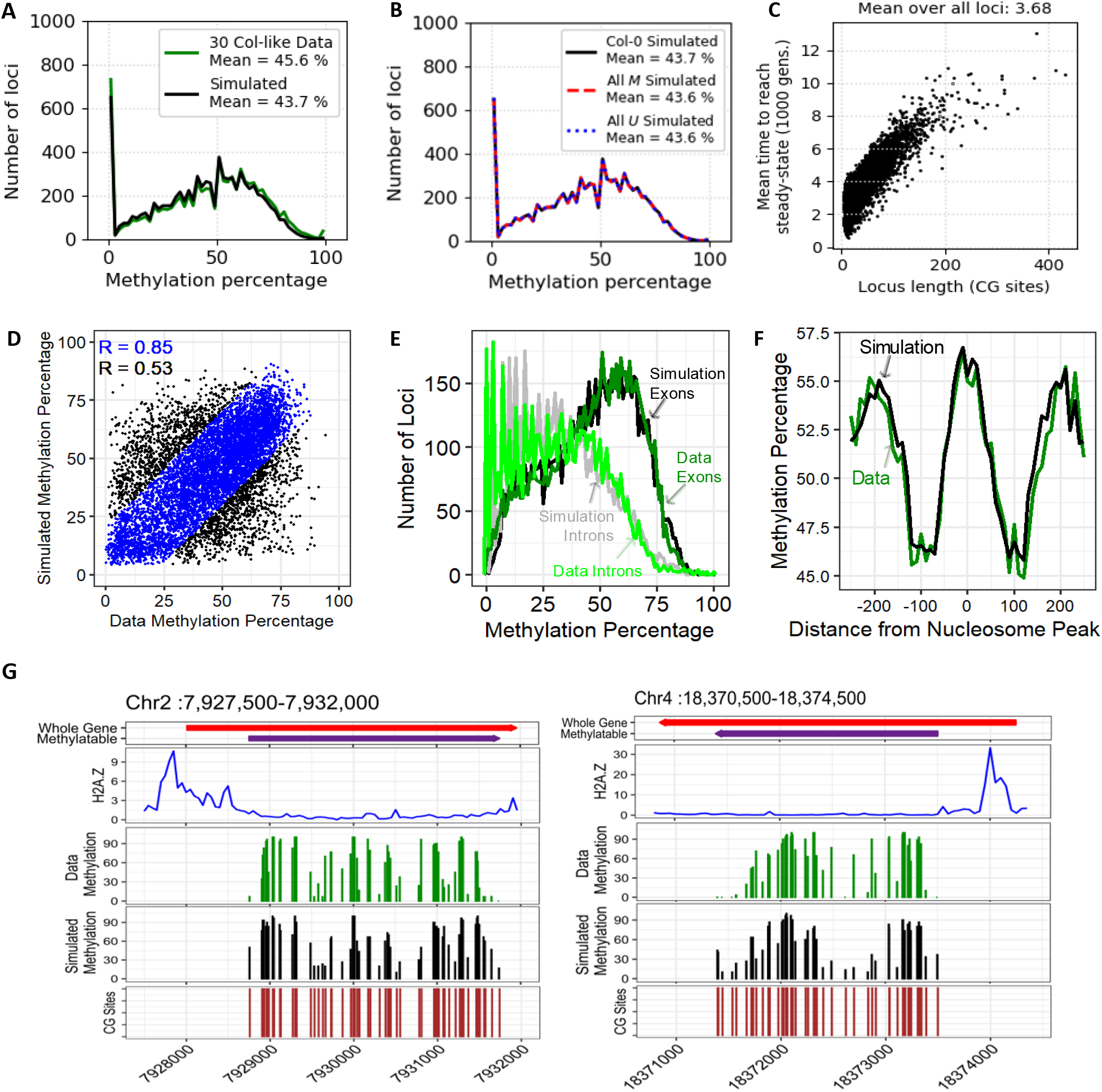
A mathematical model accurately predicts steady state gbM patterns. **A)** Histogram showing distribution of methylation levels over methylatable regions for high coverage Col-like accessions (N=30, green) and from steady-state full model simulations (black; normalized for 30 replicates after 100,000 generations starting from experimental Col-0 state; model parameters in Table S7C). **B)** Simulated steady state mCG distribution, using three initial state choices: the experimental Col-0 methylation (black, as in **A**), completely unmethylated (blue dotted) and fully methylated (red dashed). All three simulations converge to the same steady state after 100,000 generations (normalized for 30 replicates, model parameters in Table S7C). **C)** Mean time to first reach steady state as a function of locus length. Simulated over 100,000 generations from all-*U* initial state (averaged over 740 replicates), model parameters in Table S7C. **D)** Correlation of mCG levels at individual loci between full model simulations (averaged over 740 replicates) and data (averaged over 740 Col-like accessions; R = 0.53, N=7980). Well-fitting loci (<20% difference in methylation between data and simulation) are shown in blue (R = 0.85, N=6005) while poorly captured gene regions are shown in black. Simulations run as in **C**. **E)** Modelled regions were divided into exons and introns, and average levels of methylation were calculated for each region across Col-like accessions (Data, dark green for exons and light green for introns, N=30) and simulated realizations (Simulation, black for exons and grey for introns, N=30). Simulated for 100,000 generations from all-*U* initial state, model parameters in Table S7C. **F)** Methylation levels over nucleosomes are enriched in simulated methylation patterns (black, simulated as in **E**) and in the data (green). Methylation patterns averaged over well-positioned nucleosomes as defined in (Lyons & Zilberman, 2017). **G)** Genome browser views of AT2G18220 (left) and AT4G39520 (right), with gene annotation in red and the modelled methylatable region in purple. Methylation of individual cytosines agrees well between the average of 30 Col-like accessions (Data Methylation, green) and the average of 30 model realizations (Simulated Methylation, black). Col-0 H2A.Z enrichment and positions of individual CGs within the modelled region are indicated. Simulated for 100,000 generations from all-*U* initial state, model parameters in Table S7C.

Epimutation rates can be calculated over the methylatable regions using both the data and the simulated methylation changes over 30 generations (Fig. S10D; Table S1; Table S9) and good agreement is observed. Based on this, we estimated steady state gbM distributions implied by the rate changes in *cmt3, ddcc*, *ros1* and *h2az* mutants (Fig. S10E-H; Table S10). Both *cmt3* and *ddcc* are predicted to cause only small decreases in steady-state gbM (Fig. S10E-F; Table S10), further supporting the primary role of MET1 in shaping *Arabidopsis* gbM. By far the largest change is predicted in *h2az*, with steady state mean gbM in methylatable regions increasing from 44% to 68% (Fig. S10H; Table S10). This increase is predicted to unfold over almost 2000 generations (Table S10), which explains why *h2az* mutants grown in the laboratory for a few generations show only small gbM increases (Table S2) (Coleman-Derr & Zilberman, 2012).

### Steady-state gbM is not bistable

So far, we have run our simulations starting from the experimental Col-0 methylation state. Over short timescales, such as 30 generations, this is essential, because epimutation rates are modulated by proximate mCG (Fig. 3B-F). However, *de novo* methylation occurs in completely unmethylated regions (Fig. 1C), and new gbM clusters can appear and expand over just a few generations (Fig. 1D; Fig. 2A; Fig. S3). Therefore, we tested the long-timescale behavior of the model to investigate whether it can reproduce the observed methylation patterns when run from a completely unmethylated initial state (All *U*). Indeed, the same steady-state average methylation level is reached as for the empirical Col-0 initial state, and convergence of the steady-state was confirmed by also starting from a fully methylated state (All *M*; Fig. 4B; Fig. S11A). Starting from an unmethylated state, the first mCG is gained, on average, after 340 generations (Fig. S11B) and a locus reaches steady-state in around 3,700 generations on average, but the time needed depends strongly on locus size, so that loci with few CG sites reach steady-state relatively quickly (∼2,000 generations), whereas loci with >200 CG sites can take 10,000 generations or more (Fig. 4C; Fig. S11C). Overall, the model is fully converged after around 10,000 generations (Fig. S11D). This confirms that steady-state methylation dynamics are independent of the initial methylation state: the model therefore predicts that eventual mCG patterns are not bistable.

### The model accurately predicts steady-state gbM at individual loci

We next tested the predictive power of the model using data that the model was not fitted to. Using a much larger set of unique, high-coverage Col-like accessions (N=740), we simulated from a completely unmethylated initial state for 100,000 generations (repeated 740 times to match the number of accessions; distribution of methylation levels shown in Fig. S12A). The observed and simulated methylation levels were compared on a gene-by-gene basis. The values for each region were averaged across accessions and compared with those equivalently averaged across the 740 simulated realizations. We found that simulated mCG levels within 75% of gene-regions agreed well with the data, having a Pearson’s R = 0.85 (Fig. 4D). When all gene-regions were considered, the poorly captured gene-regions reduce the correlation to R = 0.53.

It is instructive to examine how the distribution of methylation varies with average CG-site density and CG-site number for each gene-region (Fig. S12B). Mean mCG increases strongly with CG-site number and is predicted well by the model (Fig. S12B; Fig. S13A). This is consistent with longer-range interactions arising from the power-law decay of the cooperative gain interaction. As a result, longer loci have a greater overall cooperative gain rate, leading to higher mean mCG. In contrast, the data and model show only small mean mCG increases with increasing average CG density (Fig. S12B). The model does best with regions that have classical gbM traits (longer, less CG-dense genes with low H2A.Z) (Muyle et al., 2022) (Fig. S13B).

The model quantitatively predicts the experimental 30-generation gain/loss rate profiles plotted as a function of distance to the nearest uCG site (rather than the nearest mCG; Fig. S13C). To examine the spatial distribution of methylated/unmethylated sites, we grouped neighboring CG sites into pairs: either uCG-uCG, uCG-mCG or mCG-mCG. The predicted and observed distributions are in good agreement (Fig. S13D). mCG-mCG separations are greatly enriched at short distances compared to uCG-mCG or uCG-uCG (Fig. S13D). This is a further demonstration of local cooperativity, which will favor clustering of methylated sites.

Model predictions for mCG levels in exons and introns agree well with the data, with higher methylation levels observed for exons (Fig. 4E). The model also reproduces the reported gbM enrichment within nucleosomes (Fig. 4F) (Chodavarapu et al., 2010; Lyons & Zilberman, 2017). Overall, spatial mCG model predictions and observed patterns at individual targets are in good agreement (Fig. 4G; Fig. 5A; Fig. S11A). Hence, the model accurately predicts steady-state gbM patterns as well as gbM levels within methylatable regions given only CG site spacing as input. The level of agreement genome-wide is remarkable given the full model (Methods) only has 15 free parameters.

**Figure 5.**
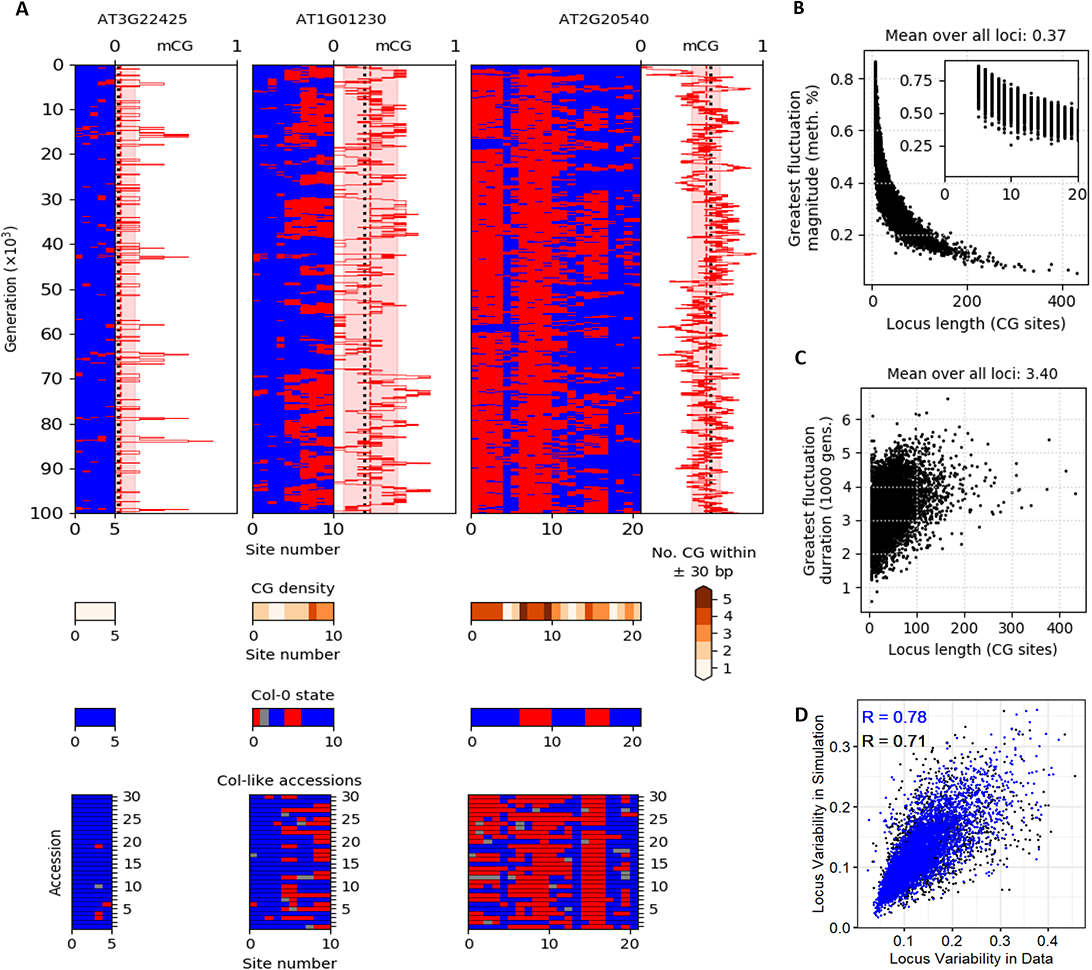
Epigenetic fluctuations explain gbM pattern variation in *Arabidopsis* populations. **A)** Top: simulated time-courses for three example loci of different size (5, 10, 21 CG sites) reaching steady-state from all-*U* initial states. Pink highlighted windows of mCG level indicate the typical magnitude of mCG fluctuations from steady state (calculated by sd of fluctuations), and the red dashed vertical line indicates the time-averaged steady-state mCG level. Black dotted vertical line shows the average mCG level of 30 Col-like accessions. Middle: local CG density over a 60 bp window centered on each CG site. Bottom: methylation patterns for Col-0 and 30 Col-like accessions. *M*-site: red; *U*-site: blue; unknown methylation status: grey; model parameters in Table S7C. **B)** Mean magnitude of greatest methylation fluctuation around mean steady-state mCG level plotted as a function of locus length. Simulation over 100,000 generations from all-*U* initial state, over 740 locus replicates: first 50,000 generations of simulation used to equilibrate, with second 50,000 generations used to measure fluctuations (model parameters in Table S7C). **C)** Mean duration (in 1000s of generations) of the greatest methylation fluctuation away from steady state, plotted as a function of locus length. Simulated as in **B**. **D)** Correlation between the standard deviation of mCG levels of individual loci within 740 Col-like accessions (Locus Variability in Data) and 740 simulated realizations (Locus Variability in Simulation; R = 0.71, N=7980). Well-fitting loci (<20% difference in methylation between data and simulation) shown in blue, as in Fig. 4D (R = 0.78, N=6005), while poorly captured gene regions shown in black. Simulations run over 100,000 generations from all-*U* initial state and averaged over 740 locus replicates (model parameters in Table S7C).

### GbM variation across natural accessions is accurately predicted by the model

Our model predicts a unique gbM steady-state for each gene, but this state is subject to substantial fluctuations, which can include completely losing and regaining methylation (Fig. 5A; Fig. S11A). Over the second half of 100,000 generation simulations (using the first half to ensure steady-state has been reached), we found that the largest absolute fractional fluctuation experienced by each locus was 0.37 on average (a change of 1 representing a transition from a fully unmethylated to a fully methylated state, or *vice versa*), and lasted 3,400 generations on average (Fig. 5B-C; Fig. S14A-B). This illustrates that gbM epigenetic fluctuations can be very large and can last a long time. This stochastic variation strongly depends on the number of CG sites in the locus (Fig. 5B), with bigger fluctuations for smaller loci, and the smallest loci remaining unmethylated for most of the simulation (Fig. 5A-C). Large clusters of CG sites maintain an overall methylated state almost indefinitely, only occasionally losing a patch of methylation (Fig. 5A-B; Fig. S11A; Fig. S14C). However, large fluctuations tend to last longer in loci with more CG sites (Fig. 5C).

Consistent with the above, loci with few CG sites have more variable gbM across natural accessions, and this is captured well by the model (Fig. S14D). The methylation levels of highly variable loci are predicted least accurately (Fig. S13B), but the variability itself is accurately predicted, with strong positive correlation between the variability of individual loci in the data and in the simulated results (R=0.71; Fig. 5D). Thus, the stochastic fluctuations of the model represent gbM variation across *Arabidopsis* accessions. The remarkable ability of our purely epigenetic model to accurately predict gbM variance across natural accessions indicates that gbM patterns in Col-like accessions (74% of assayed accessions) are primarily different stochastic realizations of an identical epigenetic inheritance process.

### Modelling outlier gbM accessions

Although most *Arabidopsis* accessions have methylomes similar to Col-0, some have global gbM levels that are considerably higher or lower (Fig. S9A) (Dubin et al., 2015; Kawakatsu et al., 2016; Pignatta et al., 2014). For example, Dör-10 has the highest overall gbM, and North Swedish accessions (NS, excluding Dör-10) have generally elevated gbM, whereas Can-0, UKID116, Cvi-0 and the group of Relicts (RL, excluding Can-0 and Cvi-0) are more sparsely methylated (Fig. S9A). The distributions of methylation levels (calculated over our methylatable-region annotation) for these outlier accessions are shown in Fig. S15A. To explore the extent to which our model can reproduce these methylation levels, we focused on two model parameters: the strength of the cooperative interaction (r*) and the level of spontaneous *de novo* activity (r0^+^; Fig. S15B-C). Adjusting only the cooperativity strength (Fig. S15B) produces reasonable fits in all cases (Fig. 6A-D), though sometimes significantly underestimates the number of fully unmethylated loci, represented by the height of the spike at the origin (Fig. 6D). In comparison, adjusting only the spontaneous *de novo* activity performs similarly for all but the most sparsely methylated accessions (Fig. 6A-D; Fig. S15C). The best fit to Can-0, UKID116 and Cvi-0, however, is found by adjusting both the cooperativity and the spontaneous *de novo* strength (Fig. 6D). Overall, as the cooperative interaction is non-linear, smaller changes in its strength are required to alter the methylation level. Our results indicate that relatively small changes to the gbM system can accommodate the entire range of gbM levels observed across the *Arabidopsis* population.

**Figure 6.**
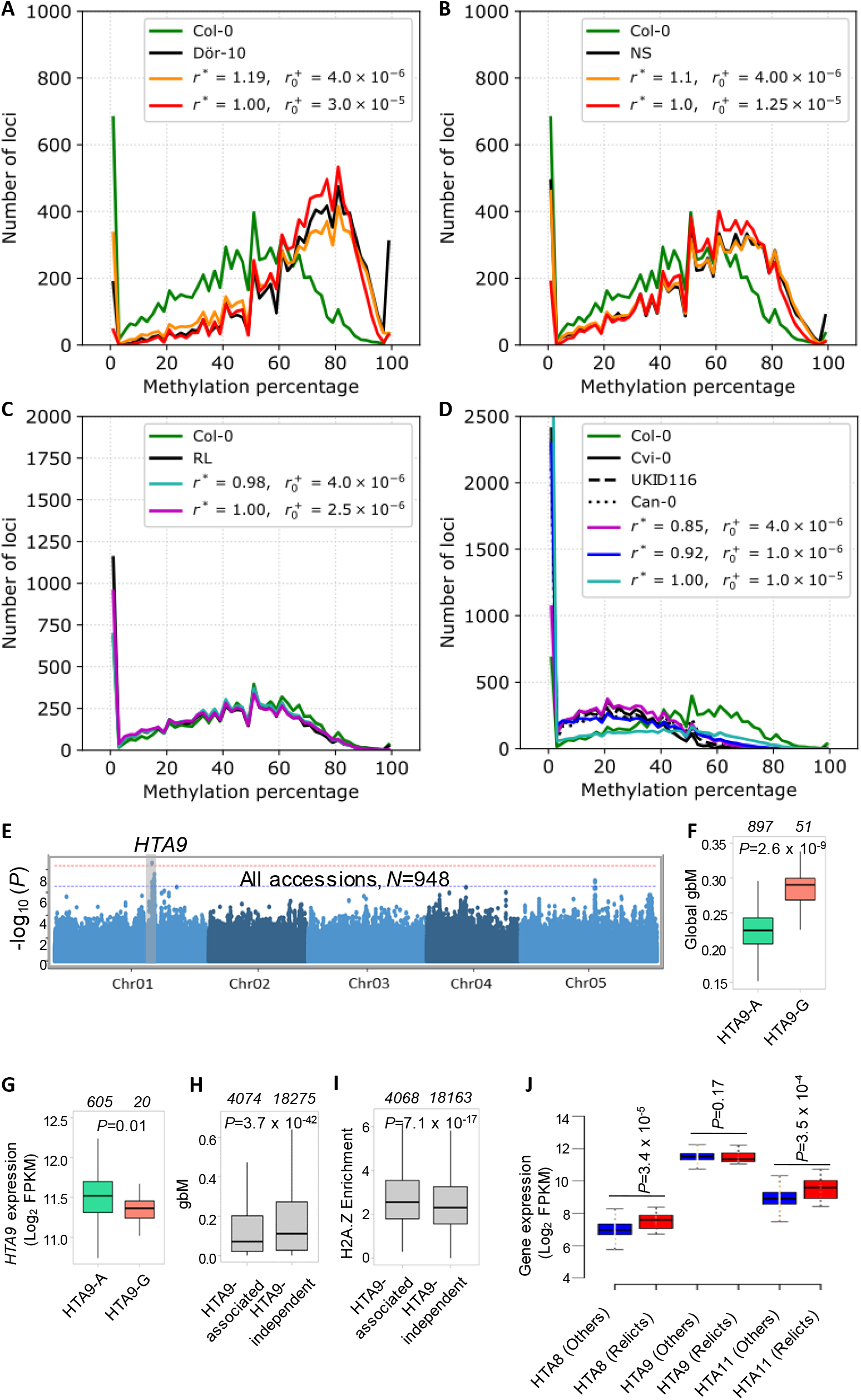
H2A.Z genetic variation is associated with global gbM variation in the population. **A)** Distribution of mCG levels for loci within Dör-10 (black) and simulated steady states for adjusted co-operative interaction strength (*r*^∗^, orange) or adjusted *de novo* activity level (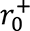, red). Col-0 distribution also included (green). Simulations run for 100,000 generations starting from all-*U* initial state, normalized for 30 realizations. Unspecified model parameters as in Table S7C. **B)** Distribution of mCG levels for loci within Northern Swedish accessions (NS, black) and simulated steady states for adjusted co-operative interaction strength (*r*^∗^, orange) or adjusted *de novo* activity level (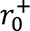, red). Col-0 distribution also included (green). Simulations as in **A**. **C)** Distribution of mCG levels for loci within Relict accessions (RL, black) and simulated steady states for adjusted co-operative interaction strength (*r*^∗^, turquoise) or adjusted *de novo* activity level (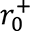, purple). Col-0 distribution also included (green). Simulations as in **A**. **D)** Distribution of mCG levels for loci within Cvi-0 (solid black), UKID116 (dashed), Can-0 (dotted) and simulated steady states for adjusted co-operative interaction strength (*r*^∗^, purple) or both adjusted co-operative interaction strength and *de novo* activity level (*r*^∗^and 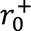, blue). Adjusting only the *de novo* activity level (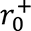, turquoise) and the Col-0 distribution are also included (green). Simulations as in **A**. **E)** *HTA9* was identified as a significant quantitative trait locus at both False Discovery Rate >0.05 (dashed blue line) and the Bonferroni threshold (dashed red line) from GWA mapping using global gbM levels as the phenotype. Y-axis indicates the –log10 of the p-values of association between SNPs and global gbM variation. **F)** Accessions harboring minor *HTA9-G* allele exhibit significantly enhanced global gbM levels compared to those with *HTA9-A* allele (p = 2.6×10^-9^, MLM GWA mapping). Number of accessions in each group indicated at top. **G)** Expression of *HTA9* is lower in accessions with the *HTA9-G* allele (p = 0.01, Wilcoxon rank sum test). FPKM=Fragments per kilobase of transcript per million mapped reads. Number of accessions in each group indicated at top. **H)** Genes whose gbM levels are associated with *HTA9* SNPs (*HTA9*-associated) have significantly lower gbM than *HTA9*-independent genes (p = 3.7×10^-42^, two-tailed t-test). Number of accessions in each group indicated at top. **I)** Levels of H2A.Z enrichment in Col-0 are significantly higher in *HTA9*-associated genes (p = 7.1×10^-17^, two-tailed t-test). Number of genes indicated at top. **J)** Expression of *HTA8*, *HTA9*, and *HTA11* in relicts (N=20) and other accessions (N=595). p-values correspond to Wilcoxon rank sum test.

### H2A.Z polymorphism is associated with high global gbM

The necessity to modify model parameters to reproduce the methylation patterns of outlier accessions implies genetic differences from Col-like accessions in factors that mediate gbM. To reveal such factors, we performed GWA analysis using the global gbM levels of accessions within the 1001 methylomes data. At the most stringent Bonferroni threshold, this identified one QTL (Fig. 6E; Fig. S16A-D), which contains *HTA9,* one of three genes encoding the H2A.Z protein in *Arabidopsis* (Yi et al., 2006). The minor HTA9 allele (HTA9-G) is found exclusively in the north, mainly in northern Sweden (Fig. S16E; Table S6), a region where accessions tend to have high gbM (Dubin et al., 2015; Kawakatsu et al., 2016). Indeed, accessions harboring the HTA9-G allele exhibit greatly enhanced global gbM (Fig. 6F) and show significantly lower expression of *HTA9* (Fig. 6G). Given our finding that *h2az* mutants have elevated rates of methylation gain and reduced rates of methylation loss (Fig. 1A-C; Fig. 2B-D; Table S1), lower expression of *HTA9* is likely responsible for high gbM in HTA9-G accessions. Dör-10, the accession with the highest gbM (Fig. S9A; Fig. S15A), harbors not only the HTA9-G allele, but also the ROS1-St and ROS3-H8 alleles that are associated with a high frequency of rarely methylated genes (Fig. 2E; Fig. S4E; Table S6). This allelic combination may account for the extraordinarily high Dör-10 gbM levels.

In addition to global gbM, we performed GWA analysis using gbM levels of individual genes. We detected *HTA9* association with gbM variation in 4074 genes (Fig. S16F-H; Table S11). These genes have relatively low gbM across accessions (Fig. 6H) and exhibit significantly enhanced deposition of H2A.Z in Col-0 (Fig. 6I). Therefore, genes with high H2A.Z appear to be more sensitive to the effects of *HTA9* genetic variation.

*HTA9* expression is not significantly different between low-gbM Relicts and other accessions (Fig. 6J). However, *HTA8* and *HTA11*, the other two *Arabidopsis* H2A.Z genes, have >40% higher expression in Relicts (Fig. 6J), which may partly account for the low gbM in these accessions. Taken together, our results indicate that H2A.Z variation is an important driver of global gbM variation in natural *Arabidopsis* populations.

## Discussion

Our results demonstrate that gbM epigenetic dynamics are dominated by MET1. In addition to its canonical semiconservative maintenance activity (Tirot et al., 2021), MET1 has *de novo* activity that is stimulated by proximate mCG (Fig. 1; Fig. 3C). Proximate mCG also boosts the efficiency of MET1-mediated maintenance (Fig. 3B). A balance of *de novo* and maintenance not only maintains existing gbM but initiates and expands new gbM clusters (Fig. 1C-D). Rates of mCG loss and gain, including gain rates in completely unmethylated sequences, are barely altered either in our *cmt3* and *cmt2cmt3* data, or in published *cmt3* and *suv4/5/6* data (Fig. 1A-C; Fig. S2F). This argues against the proposal that CMT3 plays a central role in gbM establishment (Bewick & Schmitz, 2017). Although a role for CMT3 in gbM maintenance is supported by the very low gbM levels observed in plant species that lack CMT3 (Bewick et al., 2016), our results indicate that *Arabidopsis cmt3* mutants should have only very modestly altered steady state gbM (Fig. S10E; Table S10). This suggests either that CMT3 has a stronger influence on gbM in other plant species, or that the correlation between CMT3 loss and low gbM is not causal. It is also possible that the effect of CMT3 loss on gbM dynamics increases over time, potentially due to global epigenetic alterations involving TE methylation that take many generations to unfold.

Our results also show that MET1 activity within genes is regulated by ROS1 and H2A.Z (Fig. 1A-C), but the effects of these factors are different. ROS1 activity is strongest in UM genes that are rarely methylated in the population, weakens as population gbM frequency rises, and is weakest in gbM genes (Fig. 1A-C; Fig. 2B-D). This suggests that ROS1 (likely in collaboration with other factors) determines the population gbM frequency of each gene. H2A.Z also preferentially affects mCG gains in UM sequences (Fig. 1A-C), likely because these have high H2A.Z levels (Coleman-Derr & Zilberman, 2012; Zilberman et al., 2008). However, H2A.Z reduces gains in all sequences we analyzed, including gbM segments (Fig. 1A-C; Fig. 2B-D), and therefore appears to broadly suppress gbM. This suggests that gbM may only stably exist in sequences with low H2A.Z, but because DNA methylation reduces H2A.Z abundance (Zilberman et al., 2008), H2A.Z and gbM may have a complex, dynamic relationship. This complexity is not included in the model, where we treat H2A.Z as a static background that constrains gbM regions. However, this is one of several factors that could dynamically alter on long timescales, including mCG-induced mutation of CG sites (Xia et al., 2012). The observation that H2A.Z affects mCG epimutation rates also relates to our earlier conclusion that H2A.Z has a global but generally small effect on DNA methylation levels (Coleman-Derr & Zilberman, 2012). The *h2az* epimutation rate changes are such that many generations would be required to reach a new methylation steady state (Table S10), so that laboratory measurements after only a few generations show small effects in *h2az* mutants, whereas our population genetic analyses indicate that even a modest change in H2A.Z expression can cause a major long-term alteration of the gbM landscape (Fig. 6E-J). Indeed, the long timescales involved in these processes underline the importance of modeling, where these timescales are easily accessible.

The gbM epigenetic dynamics we describe unify establishment and maintenance, and differ from other models of DNA methylation in our analysis of both processes, as well as in our prediction of a unique steady-state for each sequence (i.e., absence of bistability). Our model does not contain distinct establishment and maintenance phases: the apparent distinction between the two is driven by the sensitivity of *de novo* methylation and semiconservative maintenance to proximate mCG. Without nearby mCG to stimulate cooperative effects, gbM establishment in an unmethylated region is very rare, much rarer than cooperative *de novo* mCG addition. Nonetheless, establishment and maintenance are a single, continuous process. Any level of methylation is stable over a few generations due to high maintenance fidelity and low *de novo* activity (even with cooperativity). However, this stability is ephemeral. Due to the slight imbalance between *de novo* addition and maintenance failure, the methylation level will drift towards a unique steady state (Fig. 4B; Fig. 5A) over several thousand generations (Fig. 4C). For shorter loci, the steady state is highly unstable, as they cycle through methylation establishment, maintenance, and loss (Fig. 5A). Thus, our model differs fundamentally from previous models that predict bistability (Haerter et al., 2014; Lövkvist et al., 2016; Sontag et al., 2006; Zagkos et al., 2019). As our model successfully predicts gbM features that are common across plants and animals, including enrichment in exons and nucleosomes (Fig. 4E-F), it likely generally describes MET1/Dnmt1-mediated gbM epigenetics.

Our results have important implications for the relative contributions of genetics vs. epigenetics to gbM pattern variation across different timescales. Our hypothesis that the same processes shape gbM over short and long timescales is validated by our success in predicting steady state gbM patterns using a model based on short-term epigenetic dynamics (Fig. 4; Fig. 5A). Over short timescales, such as 30 generations, our model recapitulates the observed, essentially epigenetic gbM variation (Fig. 3E-F; Fig. S10A-C). However, over thousands of generations, gbM location and steady state are determined by factors that include CG site number and density, H2A.Z, ROS1, ROS3, and likely MET1 itself, a haplotype of which is associated with high global gbM in *Arabidopsis* (H9; Fig. S16I; Table S6). Thus, the overall gbM landscape is genetically determined. Nonetheless, the gbM steady state undergoes continuous stochastic fluctuations at each gene. The fluctuations can be large and can last for thousands of generations (Fig. 5A-C). These intrinsic fluctuations are epigenetically inherited, and they closely match the gbM differences within the *Arabidopsis* population (Fig. 5D). Thus, stochastic epigenetic inheritance generates most of the observed gbM variation between natural *Arabidopsis* accessions. This supports the earlier conclusion that *Arabidopsis* gbM variation is primarily epigenetic (Schmitz et al., 2013) and indicates that gene body methylation is an epigenetic genotype that can mediate phenotypic evolution in the *Arabidopsis* population (Shahzad et al., 2021).

## Materials and Methods

### Biological materials and growth conditions

Arabidopsis thaliana plants were grown under long day conditions (16 hours light, 8 hours dark), using the generational pattern described in Figure S1A. For *WT* (Col-0), *cmt2* (*cmt2-3*), *cmt3* (*cmt3-11*) and *drm1drm2* (*drm1-2drm2-2*), plants were grown for six total generations (G0-G5), in two separate branching lineages. For *cmt2cmt3* (*cmt2-4cmt3-11*), *h2az* (*hta8-1hta9-1hta11-1*) and *ros1* (*ros1-3*) genotypes, plants were grown for 4 total generations (G0-G3), again in two separate branching lineages. For *ddcc* plants (*drm1-2drm2-2cmt2-4cmt3-11)*, three separate G0 plants were grown for four generations (Lines A-C), each in two branching lineages.

### Leaf genomic DNA isolation, library preparation, bisulfite conversion, and sequencing

Genomic DNA (gDNA) was extracted from 1-month-old *Arabidopsis thaliana* rosette leaves with the DNeasy plant mini kit (Qiagen, cat. no. 69104) per the manufacturer’s instructions. Libraries were prepared from roughly 500 ng of purified gDNA that was sheared to approximately 400 bp on a Diagenode Bioruptor Pico water bath sonicator. Libraries were produced with the Ultra II DNA Library prep kit according to manufacturer’s instructions (New England Biolabs, cat. no. E7645L). Bisulfite conversion of DNA was carried out according to manufacturer’s protocol (Zymo, EZ DNA Methylation Lightning Kit, cat. no. D5046). DNA was converted twice to ensure complete bisulfite conversion of unmethylated cytosine. NEBNext Multiplex Oligos with methylated adaptors (cat. no. E7535L) were used for generating multiplexed libraries during PCR amplification of libraries. Sequencing was carried out as single-end 75 bp reads on Illumina NextSeq 550 at the John Innes Centre.

### Sequence Alignments and Segmentation

Bisulfite sequence reads were accessed for the mutation accumulation lines (MAL) (Becker et al., 2011; Schmitz et al., 2011), 1001 methylomes (Kawakatsu et al., 2016), and Hazarika (Hazarika et al., 2022) experiments from the Sequence Read Archive (SRA). In-house, Hazarika and published MAL sequence reads were aligned to the Arabidopsis TAIR10 genome reference sequence (Lamesche et al., 2012), using an in-house alignment pipeline as previously described (Lyons & Zilberman, 2017). 1001 Methylomes sequence reads were aligned using BSMAP 2.90 (Xi & Li, 2009) and known SNPs and indels (The 1001 Genomes Consortium, 2016) were masked. Genes and transposons were annotated using the Araport11 annotation (Cheng et al., 2017). Methylomes were segmented into Unmethylated, gbM and TE-like methylated segments as previously described (Choi et al., 2020)(Table S12). For the Col-0 data, a segmentation model was created by segmenting the combined reads from all of the Generation 3 samples in data from (Schmitz et al., 2011). Genes were defined as gbM genes if they both contained a gbM segment and any TE-like segment was both less than a quarter the length of the gbM segment and smaller than 3 CG sites (N=14,581, Table S13). Genes were defined as unmethylated genes if they contained no gbM segment or TE-like segment and did not overlap with any gene that did (N=12,045).

### Methylation Calling

Methylation status of individual CG sites was called by comparing the counts of aligned reads indicating methylated and unmethylated status at the site. Fisher’s Exact test (p<0.005) was used to determine whether there was sufficient read coverage at the site to distinguish the site from a fully unmethylated site with an error rate similar to the methylation rate observed in the chloroplast of the sample in question (as an estimate of bisulfite conversion inefficiency), or from a fully methylated site with a similar error rate. For sites where these tests indicated coverage was sufficient, a binomial test (p<0.05) was used to identify sites with significantly more methylated reads than expected from an unmethylated site. These sites were considered to be methylated. Sites which did not have significantly more methylated reads than an unmethylated site but did have sufficient reads to pass the Fisher’s exact test were considered to be unmethylated.

### Calculation of Gain and Loss Rates

For the MAL, sites with significantly more methylated reads than would be expected for an unmethylated site, but with less than 45% reads methylated, were classified as partially methylated, generally treated as missing data and assumed to consist of somatic changes and heterozygously methylated sites. A consensus methylation call was made for each site based on concordance between methylation calls for the two sibling replicates representing each line. A ‘parental consensus’ methylation state was calculated for each CG site by taking the majority methylation state among those generation 3 lines where sibling replicates agreed. For MAL, methylation state changes over 30 generations were identified by comparing parental consensus states with the states in individual lines where sibling replicates agreed, and masking known DMRs (Schmitz et al., 2011) so that we could focus on spontaneous changes at individual sites. One MA line from the Becker et al. dataset (Line 79) was an outlier in comparison to other MAL results and was therefore excluded. Over the course of 30 generations, most changes will have segregated out into the homozygously methylated or homozygously unmethylated state (see below). This, and the exclusion of sites that are heterozygously methylated at the beginning of the experiment, using the partial methylation cutoff and the ‘parental consensus’ means that MAL gains reflect those that change over the course of 30 generations from homozygously unmethylated to homozygously methylated and losses reflect those that change over thirty generation from homozygously methylated to homozygously unmethylated. The number of gains and losses was then divided by 30 to produce a per generation rate.

For our bisulfite sequencing data, sites with significantly more methylated reads than would be expected for an unmethylated site, but with less than 25% reads methylated, were classified as partially methylated, and generally treated as missing data, again to exclude somatic methylation gain and heterozygously methylated sites. Changes over individual generations were identified, using a methylation level cutoff of 70% in the parent (for losses) and offspring (for gains) to exclude segregating heterozygotes. To reduce noise, and further exclude segregating heterozygously methylated sites, only sites where parental and offspring statuses were maintained in the six closest relatives were considered. The result of this is that gains reflect those which change over one generation from homozygously unmethylated to homozygously methylated and losses reflect those which change over one generation from homozygously methylated to homozygously unmethylated. This method will not capture all changes, but will underestimate the rate by a factor of two. To illustrate, a gain represents a site that begins as homozygously unmethylated (UU) in Generation 0. Occurrence of a gain will result in the site being heterozygously methylated (UM) at Generation 1 (we assume that changes happen on only one chromosome due to the low rate of epimutation). Mendelian segregation predicts three potential outcomes in Generation 2: 25% of all UM sites will return to being UU, and will not result in a gain in methylation. 25% of all UM sites will become MM, fixing the gain in the population. Using our method, only these changes will be detected. The final 50% of UM sites will remain UM in Generation 2. These gains will not be detected, as their methylation level in Generation 2 is not expected to be above 70%, the cutoff used here. Half of these UM sites will eventually, over multiple generations, become fixed as MM and therefore result in methylation gain. However, our method does not detect these gains as they are not yet fixed in the population, and we therefore multiply by two the number of gains that we do detect to include these sites. Individual comparisons between plants with rates that were strong outliers (values +/- 1.5 S.D. from the average) were excluded.

For all methods, rates of change per site were estimated by dividing inferred gains/losses by the number of *U* calls/*M* calls in the previous generation. The resulting rates per generation were converted to per cell cycle by a fixed estimate of 34 germ cell cycles per generation (Watson et al., 2016).

### Population gbM frequency

Population gbM frequency of each gene was obtained from (Shahzad et al., 2021) and were calculated as described there. Briefly, the population gbM frequency represents the number of accessions having gbM at a given gene as a percentage of the total available calls of gbM, teM or unmethylated across 948 *Arabidopsis* accessions.

### Choice of accessions for modelling

Col-like accessions were defined as accessions where the proportion of called sites called as M, in segments annotated as gbM in Col-0 (Schmitz parental consensus data set), is within 1 SD of the mean among all samples (N=891). Accessions’ global mean gbM was calculated by identifying the region of broadly gbM-methylatable genome space, defined as regions that are covered by gbM segments in at least 5 *Arabidopsis* accessions, and counting CG sites in that space called as methylated in the accession by binomial test, as a proportion of sites called as methylated or unmethylated. Individual accessions’ global mean gbM was calculated excluding any portion of the broadly gbM-methylatable space that is overlapped by a teM segment in the accession in question.

To fit the model, accessions with more than 60% of CG sites called as either methylated or unmethylated (adequate coverage) were divided into groups based on their global mean gbM. The 10 most hypo- and hyper-gbM accessions (∼1%) were identified as potential outliers and removed from the broader group. The remainder of accessions were divided into deciles by global mean gbM. The 10 accessions with highest proportion of CG sites called as methylated or unmethylated (best coverage) in each of the three deciles closest to Col-0 (Deciles 3, 4 and 5) were chosen to determine model parameters (Data_Fit, N=30)

For a wider comparison of the model, simulations were compared to all unique, high-coverage Col-like accessions. Duplicate samples of individual accessions were removed (N=798). Accessions with less than 50% of CG sites called as either methylated or unmethylated were considered to have inadequate coverage and were removed, resulting in a final accession set for modelling of N=740.

To analyze accessions with non-Col-like levels of methylation, two groups were chosen: Northern Swedish accessions (NS), which have elevated gbM, and Relict (RL) accessions, which have unusually low levels of gbM. Dör-10, a Northern Swedish accession which has the highest known level of gbM was analyzed separately, as were two Relict accessions (Can-0 and Cvi-0), which have extremely low levels of gbM, and a third accession (UKID116), which has similarly low levels of gbM.

### Methylatable Gene Regions

Genes for modelling were selected from the Araport 11 annotation, annotated as protein coding – (N=27,473). Genes that contained TE-like methylation in greater than 1% of accessions were discarded to retain non-TE genes only (N=19,082). The segments chosen for modelling comprise the region in each gene between the first and last CG sites overlapped by gbM segments spanning at least 3 CG sites, in at least 5% of the accessions from the 891 which constitute the Col-0 like data set, resulting in the regions that were broadly methylatable (N=13,138). These segments were further trimmed to remove the ends of segments where H2A.Z ChIP-Seq signal, smoothed over 5 adjacent 50bp bins, was above 1.2. H2A.Z ChIP data was obtained from (Coleman-Derr & Zilberman, 2012). Segments were also removed if the mean H2A.Z ChIP-Seq signal along the remaining segment exceeded 1.2, resulting in the H2A.Z-low methylatable annotation (N=8,843). Our modelling method considers methylation levels in each locus in isolation of the surrounding DNA and therefore overlapping genes (where gbM in one gene may affect gbM in the other, overlapping gene) cannot be appropriately modelled. To prevent this, if segments overlapped by more than 20%, the smaller segment was discarded (N=416), as were any gene regions containing fewer than 5 CG sites (N= 447). This left us with 7980 genes in our final modelling dataset (Table S8).

### Analysis of sparsely methylated regions

Co-ordinates of sparsely methylated regions within gbM genes (SPMRs) and the gene IDs for the gbM genes concerned (Hazarika gbM genes) were downloaded from (Hazarika et al., 2022). Regions within the Hazarika gbM genes that were not SPMRs were defined as non-SPMRs. Non-SPMRs were bimodal with respect to mCG and so were partitioned into methylated (M_non_SPMRs) and unmethylated (U_non_SPMRs) (Fig. S2A-C). 91% of M_non_SPMR CG sites and 92% of SPMR CG sites fall within our gbM segments. Epimutation rates in both published MAL data and our bisulfite sequencing data were calculated in SPMRs, M_non_SPMRs and U_non_SPMRs (Fig. S2D; Table S1).

### Reanalysis of Hazarika et al. methylation data

Bisulfite sequencing data was downloaded from Hazarika et al., 2022 and methylation levels were calculated by averaging the percentage of C reads/total reads at each site over each genomic region. Samples are of highly variable coverage (4-83X). In the original analysis, in order to make single site methylation calls in all samples, methylation states of individual cytosines were imputed based on the methylation status of nearby cytosines (Taudt et al., 2018). This method may not be appropriate to make single site mCG calls in sparsely methylated genes, especially to subsequently identify rare epimutations within an otherwise unchanged methylation pattern of neighboring sites. We therefore excluded samples of coverage <10X (WT Line 1 Generation 11, *suvh4/5/6* Line 4 Generations 5 and 13, *suvh4/5/6* Line 8 Generation 8 and 9) and assigned methylation status without imputation in all remaining samples. Sites with significantly more methylated reads than would be expected for an unmethylated site, but with less than 25% reads methylated, were classified as partially methylated, and generally treated as missing data. A ‘parental consensus’ methylation state was calculated for each CG site by identifying shared *M* or *U* calls in the three samples at the earliest generations for each genotype. Methylation state changes were identified by comparing parental consensus states with the states in individual samples and converted to per generation rates depending on the number of generations since the parental consensus. Rates were calculated in different genomic regions.

### Genome-wide association mapping

Three types of genome-wide association (GWA) analyses were performed to identify single nucleotide polymorphisms (SNPs) associated with the natural variation of gbM in *Arabidopsis* accessions. Genome-wide average gbM levels of accessions were used to identify the genetic factors influencing global gbM. Global gbM levels are influenced by sequencing coverage (Fig. S16A) and thus analyses were carried out using multiple cutoffs for sequencing coverage.

Additionally, gbM levels of individual genes were used for GWA analysis to identify SNPs linked with local gbM variation. Furthermore, gbM frequency in rarely methylated genes (RMGs) was used for GWA analysis. RMGs are defined as genes with gbM in <20% of accessions and which have mean H2A.Z ChIP-seq signal <4 in Col-0 gene bodies. GWA mapping was performed using 1001 genomes SNP data (The 1001 Genomes Consortium, 2016) with an accelerated mixed model (AMM) (Seren et al., 2012) implemented in PyGWAS: Python library for running GWAS (version 1.7.4). SNPs with Minor Allele Frequency (MAF) >5% in the population were considered. 0.05 False Discovery Rate (FDR) correction (Benjamini & Hochberg, 1995) was implemented to account for multiple tests and identify SNPs associated with gbM variation.

### Haplotype analyses

ROS1 and ROS3 nucleotide sequences of *Arabidopsis* accessions were retrieved from http://signal.salk.edu/atg1001/3.0/gebrowser.php (The 1001 Genomes Consortium, 2016). Sequences were translated with the EMBOSS Transeq pipeline, and aligned using MEGA5 (Tamura et al., 2011). Amino acid polymorphisms were identified and accessions identical (100%) for predicted full length protein sequences were classified as a haplogroup. Haplogroups comprising less than 15 accessions were discarded from association analyses to have a reasonable number within each haplogroup. The ROS1 stop codon group includes all the accessions with a premature stop codon irrespective of the stop codon position and predicted protein amino acid sequence identity. Associations between ROS1 and ROS3 haplogroups and gbM variation were examined using the linear model ANOVA.

### Modelling gene body methylation dynamics

Methylation dynamics are modelled over both short timescales (30 plant generations) and long timescales (100,000 plant generations, corresponding to over three million cell cycles). Despite the remarkably high fidelity of the maintenance pathway, over these long timescales maintenance failure events will accumulate. We study the role of cooperative *de novo* and cooperative maintenance pathways in generating stable methylation dynamics consistent with the experimentally observed methylation patterns. The term ‘cooperative’ is used to denote, for example, that the probability of a specific unmethylated CG site (uCG) gaining methylation is enhanced by the presence of other nearby methylated sites (mCG).

A single copy of each diploid chromosome is modelled through multiple cell cycles, assuming that the methylation dynamics are unaffected by whether that chromosome is currently in a diploid or haploid cell environment. Below we discuss in detail how this choice relates to the experimental setups. Furthermore, we model only the CG sites and assume that the methylation dynamics of each gene are independent from all others so that each gene can be modelled individually. Genes (or methylatable-regions) consisting of only four CG-sites or fewer are not simulated. Subtleties occurring when genes overlap are discussed above in the section “Methylatable Gene Regions”. The methylation dynamics are simulated stochastically using the direct Gillespie algorithm (Gillespie, 1977), as described below, with the cooperativity implemented using an independent interaction between each target uCG and every mCG. All timescales and rates are defined relative to the cell cycle duration (here taken to be the time between successive DNA replication events).

### Construction of Two-State Model

The model does not keep track of the individual methylation status of the top and bottom strand cytosines of each CG site. Instead, a single CG site is defined to be in one of three states: fully-methylated (*M*), hemi-methylated (*H*) or un-methylated (*U*). Initially we assume that there is no active demethylation occurring in gene bodies (e.g., through the DNA glycosylase pathways). We later revisit this assumption, however, in light of the *ros1* dataset. The possible methylation gain and loss transitions between the three states are illustrated and defined in Fig. S6A.

During replication, all *M* sites are converted to *H* sites. On average, we assume that half of the pre-existing *H* sites will be methylated on the stand of DNA corresponding to the lineage that we follow; these therefore remain as *H*, whereas the remainder of the pre-existing *H* sites transform to *U* sites.

It is known that the MET1 maintenance pathway is extremely efficient (Song et al., 2012), a result recapitulated by our analysis. In the analysis below, we assume that MET1-mediated re-methylation is not only highly efficient but occurs rapidly after replication on a timescale much faster than the cell cycle duration. This assumption simplifies our analysis. If, however, maintenance is slower (but still highly efficient within a cell cycle), our conclusions are nevertheless unchanged. We first define a maintenance failure rate, *f*, which is the probability per cell cycle that an *H* site generated during replication is not converted to an *M*. This gives a remethylation maintenance rate

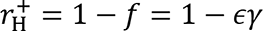

with 0 < *ε* ≪ 1, and where 0 < *γ* < 1 is a further cooperative ‘suppression-factor’, which reduces the chance of a maintenance failure occurring if there are existing *M* sites nearby (see below). In the limit of *γ* → 1, *ε* is the background maintenance failure rate that a single isolated *M* site immediately before replication, in an otherwise completely unmethylated gene, is not fully remethylated in the period between successive DNA replication events.

The *de novo* methylation of a *U* site, 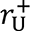 ≪ 1, is composed of both a spontaneous and a cooperative pathway as discussed below. As expected, in the experimentally relevant region of parameter-space (i.e., very efficient maintenance by MET1, *ε* ≪ 1), *H* sites only exist transiently, up to one cell cycle, before being resolved to *U* or *M*. As a result, the intermediate state, *H*, can be integrated out to create a two-state model consisting of only *U* and *M* states with effective direct transitions between these two states. This simplification was confirmed to have negligible effect on the simulation results by first implementing the three-state model described above using a Gillespie algorithm that was interrupted at the end of every cell cycle to explicitly simulate every replication event. In the experimentally applicable region of parameter-space, the three-state and two-state models produced almost identical methylation dynamics. Crucially, however, as replication is no longer explicitly simulated for the two-state model, a single Gillespie time increment can span multiple cell cycles, speeding up the simulation by several orders of magnitude. This speed up was helpful in simulating methylation dynamics over very long time periods of 10^5^ plant generations.

In the two-state model, the effective *M* → *U* loss-rate, *r*^−^, to first order is given by:

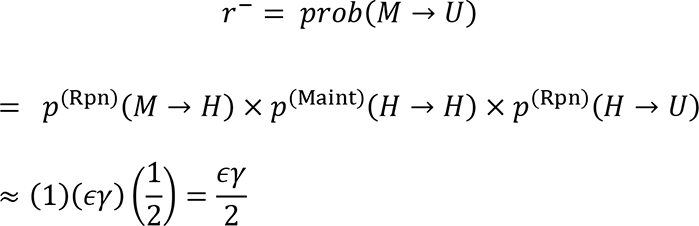

where *p*^(Rpn)^ represents the probability of the specified event occurring at replication, and *p*^(Maint)^ the probability of the specified event occurring during maintenance. Similarly, the effective gain-rate, to first order, is given by:

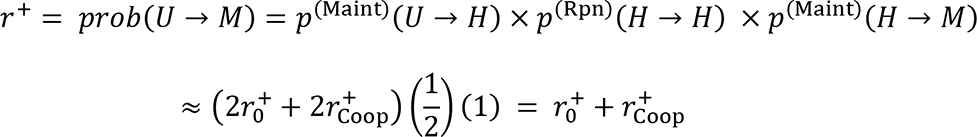

where 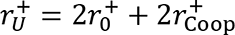 is decomposed into two components. Firstly, a constant spontaneous *de novo* gain-rate, 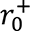, which is assumed to be a uniform background across all CG sites. Secondly, 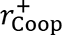 represents the cooperative gain rate. We discuss the precise implementation of this contribution in the next section. The explicit factor of 2 in the parameterization of 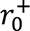 and 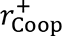 is included to account for the fact that in each *U* site there are two possible unmethylated cytosine targets for the pathways to act on. Only the first order contributions to the effective rates are included, as self-consistently, we find that the fitted values for 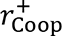, 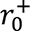 and *ε* per cell cycle are all several orders of magnitude smaller than one.

In our previous work modelling decay of methylation levels at transposable elements (TEs) in various *A. thaliana* mutants (Lyons et al., 2022), we assumed a constant methylation gain and loss rate per cell cycle without explicit cooperativity, despite the rapidly falling methylation level. In light of our current work, it seems likely that both cooperative *de novo* and cooperative maintenance will also shape MET1 activity in TEs. The previously calculated rates for TEs in (Lyons et al., 2022) therefore constitute effective ‘average’ methylation gain/loss rates (including cooperativity) during the decay dynamics.

### Gillespie simulation: cooperative gains

We simulate using the ‘direct’ Gillespie algorithm (Gillespie, 1977). An intermediate cooperative gain propensity, 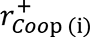, is calculated for every pair of *U* and *M* sites (*UM*-pair) in the current gene-region. The functional form of 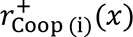, depends on *x*, the base-pair separation between the two CG-sites in the *UM*-pair and is discussed in a later section. The cooperative gain propensity, 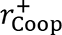, for a given *U* site is then found by summing all the individual contributions from each of the *UM*-pairs for that particular *U* site: 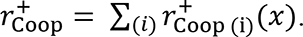. The overall gain propensity for that individual *U* site is then given by 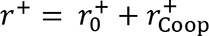 as defined in the previous section, while *M* sites are assigned *r*^+^ = 0. The total gain propensity for the gene-region, 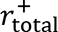, is the sum over all the individual site-propensities. Similarly, a loss propensity, *r*^−^, is calculated for every individual *M* site, as described in detail in the next section, along with *r*^−^ = 0 for *U* sites. The individual-site loss propensities are also summed to give 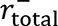 and finally the total propensity for the entire gene-region is defined as 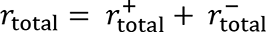.

At a time, *t*, the next event (i.e., gain or loss) will occur at time: *t* + Δ*t*, where Δ*t* = 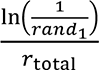 and *rand*_1_ is a uniformly distributed random number between 0 and 1. The site to be updated at this time is found using the propensity threshold: *r*_total_ ∗ *rand*_2_, where *rand*_2_ is a second random number from the same distribution. The smallest individual site-propensity for which the cumulative sum of site-propensities, up to and including that site-propensity, exceeds the threshold determines the identity of the next site to be updated (i.e., gain/loss of methylation) at time *t* + Δ*t*.

We assume that there are no cooperative interactions between CG sites in different genes. For a *UM*-pair separated by *x* = 300 bp, the cooperative gain amplitude is over a factor of 1000 smaller than its maximum value, therefore, this assumption is unlikely to have a significant impact on the simulated dynamics.

### Gillespie simulation: cooperative maintenance

We model a cooperative maintenance pathway using a similar approach to that for the cooperative gains. We assume that the presence of other *M* sites in the locus increases the chance of a site being maintained after replication. Again, we approach this in a pair-wise fashion, this time considering all *MM*-pairs in a locus. As the bare (i.e., non-cooperative) maintenance failure rate, *ε*, is already so small, we multiply the contributions from the individual *MM*-pairs to the enhanced maintenance rate. This is to ensure that the total maintenance probability can never exceed 1 at any *M* site. Each *MM*-pair contributes a ‘suppression-factor’ to the maintenance failure rate: (1 − *β*_(i)_(*x*)), (Fig. S6F), where *x* is the base pair separation between the two sites of the *MM*-pair, and 0 < *β*_(i)_(*x*) < 1, for all *x*. The loss-rate at an individual *M* target-site is then calculated as 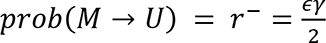, where:

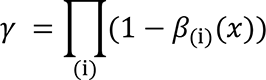

The product is taken over all *MM*-pairs that contain the specific *M* target-site being maintained. The functional form of *β*_(i)_(*x*) is discussed below and is the same for every *MM*-pair in the locus. Simulations omitting cooperative maintenance produce a uniform loss rate (equal to *ε*/2), thus failing to reproduce the suppression of methylation losses observed at short-length scales (Fig. S6E).

### Active demethylation

The *ros1* mutant data indicates that active demethylation contributes to the dynamics. One possibility is that a ROS1 pathway could renormalize the *U* → *M* transition probability so that the values of the parameters 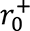 and 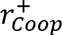 also include the action of ROS1 rapidly removing some hemi-methylation before the next replication event. In addition, ROS1 may be actively demethylating *M* sites. This is largely captured by the current model by the existing maintenance failure pathway (i.e., some proportion of the parameters *ε* and *β*_(*i*)_ could in principle arise from ROS1 action). An alternative parameterization for a ROS1 active demethylation pathway, however, might be to include a spontaneous (non-cooperative) *M* → *U* interaction (analogous to 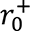 for the spontaneous gains). This is not included in the model as the current resolution of the loss-rate data is insufficient to support fitting to an additional parameter.

### Functional form of cooperative interactions

The magnitude of the cooperative gain-rate for an individual *UM*-pair, 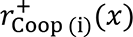, is a function of *x*, the base-pair separation between the *UM*-pair: *x* = |***x***_***U***_ − ***x***_***M***_|. We split 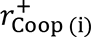 (*x*) into a primary component, 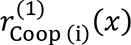, and a secondary component, 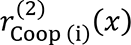, for which the interaction has the same functional-form as the primary component but with the origin translated by *x*_*nucl*_, a distance on the length scale of the nucleosome repeat length:

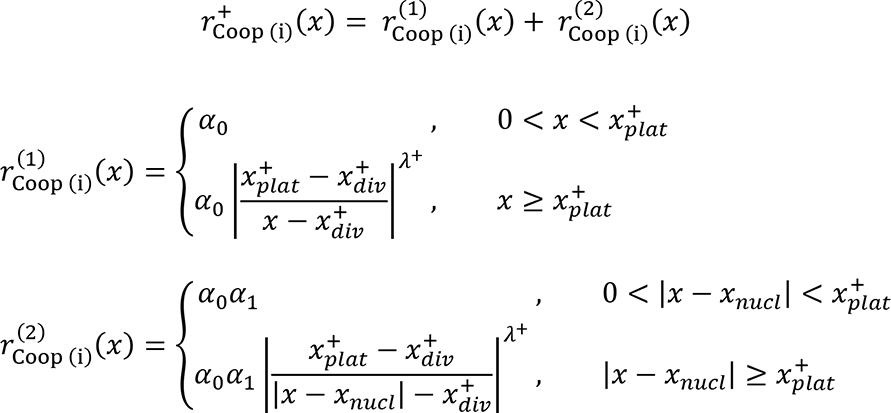

For 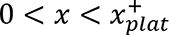, for the primary component, and for 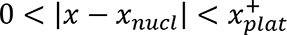 for the secondary component, both have a constant magnitude described by *α*_0_ and *α*_0_*α*_1_, respectively. This parameterisation was chosen so that the strength of the secondary component could be scaled relative that of the primary component when introducing the interaction-strength dependence on CG-density (described below). All other parameters are assumed to be equivalent for the primary and secondary components of the cooperative gain interaction. The parameter *x*_*plat*_ defines the end of this constant plateau, beyond which the interaction has a power-law decay. The location of the power-law divergence is specified by *x*_*div*_, where *x*_*div*_ < *x*_*plat*_. Finally, the power-law decay constant is specified by *λ*^+^. The secondary interaction could occur due to the 3D organization of the DNA providing an alternative, shorter, interaction route between the two CG-sites in the *UM*-pair. The overall functional form of 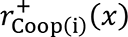 is shown in Fig. S6B.

The cooperative maintenance interaction is described analogously to the cooperative gain interaction (defined above):

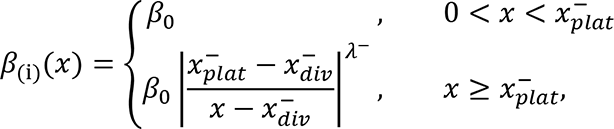

with the functional form (1 − *β*_(*i*)_(*x*)) shown in Fig. S6F. Choice of this functional form ensures convergence at long distances to the background (non-cooperative) methylation loss rate. We allow the amplitude (*β*_0_), plateau length 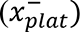, divergence location 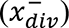 and power-law decay constant (*λ*^−^) parameters to all have values independent of those used for the cooperative gain interaction.

We chose a power-law to describe the decay of both the cooperative gain and cooperative maintenance interactions with increasing separation of the *UM*-pair. This is consistent with the linear form of the loss-rate over one order of magnitude, when plotted as a function of distance to the nearest *M*-site, using a log-log scale as shown in Fig. S6C. A linear best-fit (to the log-log transformed data 20 < *x* < 200 bp) is also shown. Note that the gradient of this linear-fit to the overall loss-rate does not provide the value of the power-law *λ*^−^ because *λ*^−^ describes the contribution to the total interaction that arises from only a single *UM*-pair. Hence, the entire shape of this interaction strength profile for a single *MM*-pair (Fig. S6F) differs from the full losses profile (Fig. 3E), due to the latter including interactions with multiple mCGs. The structure of the equivalently plotted gain-rate is more complex as there is the secondary-interaction peak at *x* ≈ 170 bp (log_10_(*x*) ≈ 2.2). In this case, we therefore, make a linear fit to the log-log transformed gain-rate over only a narrow window of 25 < *x* < 47 bp, where the primary cooperative gain pathway is dominant. The linear fit is plotted over the entire length-scale of the gain-rate data and at large x-values, beyond the range of the secondary interaction peak, is also quite consistent with the gain rate data. We note however, that as the resolution for the gain and loss rates becomes low at larger length-scales (of the order *x* > 500 bp), we cannot conclusively rule out the possibility of an exponentially decaying interaction strength, though fits to an exponential were not as good. Once again, the shape of this gains interaction strength profile for a single *UM*-pair (Fig. S6B) differs from the full gains profile (Fig. 3F), due to the latter including interactions with multiple mCGs.

For the model to correctly reproduce the observed distribution of steady-state methylation levels for gene-regions of varying length (*L*_*locus*_), and average CG-density (*ρ*_*CG*_), we had to vary the strength of the cooperative gain interaction and of the cooperative maintenance interaction, as a function of CG-density. We define *L*_*locus*_ to be the base pair distance between the first and last CG sites in the gene-region and chose the unconventional definition of 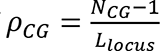, where *N*_*CG*_ is the number of CG sites in the gene-region, so that 1/*ρ*_*CG*_ is equivalent to the average CG site spacing.

To introduce as few extra parameters as possible, we used the linear scaling:

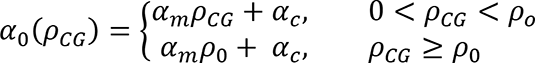

where *α*_*m*_ < 0 so that the cooperative gain interaction strength decreases with increasing CG-density, and the maximum CG-density threshold, *ρ*_0_, prevents a vanishing interaction strength. Similarly, we vary the strength of the cooperative maintenance interaction, *β*_0_, as a function of CG-density using:

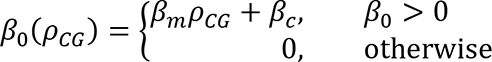

where *β*_*m*_ < 0. The cooperative maintenance strength is capped to have a minimum value of zero. Finally, when we model the outlier *Arabidopsis* accessions, we vary the strength of the cooperative gain and cooperative maintenance strengths using a single scale-factor, *r*^∗^, so that *α*_*c*_ → *r*^∗^*α*_*c*_ and *β*_*c*_ → *r*^∗^*β*_*c*_.

In summary, the final model contains four gain/loss processes: spontaneous *de novo* (one parameter); cooperative *de novo* (eight parameters); background maintenance failure (one parameter); cooperative suppression of maintenance failure (five parameters), to give 15 parameters in total.

### Calculation of diploid methylation gain and loss rates

We simulate the methylation dynamics of a single chromosome. All bisulfite sequencing used here, however, measured the methylome of diploid leaf tissue. Each of the two copies forming a chromosome pair will have had a different trajectory, alternating between a haploid and diploid environment to reach the final leaf tissue that is sequenced. Our model contains no information on whether the simulated chromosome is currently in a haploid or diploid cell, or any other details about its environment. We therefore treat the two chromosome copies as indistinguishable.

The simulated gain and loss rates are compared to two different data sets: firstly, the existing Mutation Accumulation Line experiments (MAL) (Becker et al., 2011; Schmitz et al., 2011), and secondly the *WT* dataset presented in this work. These two approaches use contrastingly shaped lineage-trees and consequently involve complementary assumptions. We convert the rates from both experimental data sets, and the simulated data, into the diploid gain/loss rate per cell cycle, i.e., for methylation gains, this is the rate that a homozygous *U* site, is converted to a homozygous *M* site.

As we now show, the diploid gain/loss rate is equivalent to the haploid gain/loss rate found by simulating a single chromosome. If we define *r* to be the haploid gain rate per plant generation for a single chromosome, then for a diploid cell, the total number of newly heterozygous-methylated sites generated by a methylation gain on one of the chromosomes in a single generation is 2*r*. In these experiments the plants are self-fertilized each generation, so for the heterozygous sites, there is a 25% chance that Mendelian segregation will result in a homozygous gain/loss, a 25% chance the gain/loss disappears so that the original homozygous state is retained, and a 50% chance that it remains heterozygous at each subsequent reproductive cycle. Eventually, therefore, there is 50% chance that a heterozygous gain is ‘fixed’ into a homozygous gain and a 50% chance it reverts to a homozygous *U* site. This factor of ½ cancels the initial factor of two and hence the diploid gain gate is equal to the haploid gain rate on a single chromosome.

We note that after the male and female cell-lineages diverge to form the reproductive tissues, the rate of producing new heterozygous gains doubles as there are now four chromosomes on which a gain could occur. For the purposes of this discussion, we chose to define the start of the generational cycle as the divergence point of the male and female lineages. Meiosis and fertilization then occur mid-way through this cycle. Neglecting the (negligible) possibility of a gain occurring at the same CG site in both the male and female lineage during a single generation, upon fertilization, there is a 50% probability that (in our followed lineage) the CG site reverts to homozygous unmethylated and a 50% probability that it persists as a heterozygous gain in the now single germline lineage. The total number of new heterozygous gains formed so far is therefore double the number of haploid gains. As there is only a single germline lineage for the remainder of the cell cycle, heterozygous gains continue to be formed at double the haploid gain rate. Consequently, the new heterozygous gains are produced at the same overall rate from before and after the male and female lineages diverges, and the first opportunity for self-fertilized Mendelian segregation then occurs part way through the following cell cycle (with losses behaving equivalently).

The mutation accumulation line experiments have a narrow and very deep tree, following 4 lines in the Schmitz et al. dataset and 8 in Becker et al. (we excluded 1 outlier line, as discussed above), all for 30 consecutive plant generations. Consider a heterozygous methylation gain occurring in the first generation. After five reproductive cycles, for example, there is a probability of only (0.5)^5^ ∼ 3% that the CG site will still be in a heterozygous state. We assume, however, that all gains and losses occurring during the observed 30 generations have fully segregated out at the point of bisulfite sequencing. With this approximation, our analysis of these 30 generation datasets directly measures the diploid gain/loss rates over 30 generations. Finally, this is converted to the per cell cycle gain/loss rate by dividing the gain/loss rate for the entire experiment by the number of intermediate generations (in this case, 30 generations) and by the number of cell divisions through the germline lineage over one generation. For the latter, we assume that there is a constant number of cell divisions of 34 per generation (Watson et al., 2016).

In reality, a proportion of the sites will remain in the heterozygous state at the end of the experiment. To analyze the Schmitz et al. and Becker et al. datasets, both the Generation 0, parental germline state, and the Generation 30, final germline state, are reconstructed from sibling offspring lines, as described previously. Defining the generational cycle to begin at the divergence of the male and female lineages is therefore consistent with aligning it to the experimental measurement of germline methylation state. Heterozygous sites in the initial or final experimental state are most likely to be classified as indeterminate methylation (*I* sites). As gains/losses are only identified at CG sites where both the initial and final methylation state can be called as either *U* or *M*, heterozygous methylated sites in the final observed state are excluded from the measured gain/loss rates. Excluding heterozygous sites that were formed prior to Generation 0 from the analysis is consistent with our assumption that all gains and losses are initiated during the 30-generation experiment. Excluding all heterozygous sites remaining in the final state at Generation 30, however, produces a slight underestimate of the true gain and loss rates.

We now estimate an upper limit to the fraction of gains lost due to the assumption that all will have fully segregated by the end of the experiment. Again, we use ‘*r*’ to represent the haploid gain rate per generation. Over the 30-generation experiment, 2*r* heterozygous gains will occur per generation. Theoretically, if we could wait until all the heterozygous gains generated during the 30 generations then fully segregated out, this would lead to 30*r* diploid gains in total. Over the 30-generation experiment, heterozygous sites formed during the *x*^th^ generation will have 30 − *x* self-fertilized Mendelian segregation opportunities. On average, therefore, 100% of the heterozygous gains occurring in the 30^th^ generation will have been missed, and 50% from the 29^th^ generation and 25% from the 28^th^ etc. The total number of heterozygous gains yet to segregate is given by the sum: 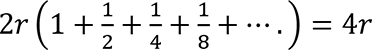. Half of these are expected, on average, to be fixed into full diploid changes, so that 2*r* gains have been uncounted. This fraction is 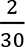 ∼ 7% of the total number of diploid gains over the entire 30 generations. An identical argument applies to the fraction of losses not accounted for. Finally, we note, however, that heterozygous sites do not explain all the CG sites identified experimentally as indeterminately methylated. The majority are likely due to somatic methylation gains. As they are concentrated close to existing methylation, we anticipate that they correspond to sites that are unmethylated in the germline but having a high gain probability. Excluding these sites, therefore results in a small underestimate of the gain rate.

In contrast, the bisulfite sequencing data presented in this work uses a much broader and shallower lineage tree. As described above, we assume that the identified methylation gains/losses are homozygous (present on both chromosomes). This rate is then doubled to give the eventual diploid gain/loss rate per generation, reflecting the additional 50% of heterozygous changes that will eventually become fixed in the changed state. Again, we convert to a rate per cell cycle.

### Simulated gain and loss rates

We fit the model described above to the experimental gain and loss rates. As we compare to our analysis of four lines of the 30-generation data set of Schmitz et al., we also simulate a single copy of each chromosome for 30 generations, assuming 34 cell cycles per generation (Watson et al., 2016). For a direct comparison to this experimental data, the simulation is also repeated for four independent replicates. The random-number seed was increased by one for every new gene-region and for each replicate. We note that as expected, simulating over a greater number of replicates reduces the fluctuations seen in the gain and loss rates, especially at long length-scales.

The simulation uses the experimental Col-0 state to assign CG sites an initial status of either *M* or *U*. To achieve the best resolution, we use the ‘Col-0 consensus state’, generated by collating high coverage bisulfite sequencing from twenty different Col-0 plants grown under consistent conditions. In this consensus state, 1.77% of sites have indeterminate (*I*) status, most likely indicating that the CG site either has a large proportion of somatic methylation changes, or that it is heterozygous in the germline. For consistency with the experimental analysis, we entirely exclude these *I* sites from the simulation for the fit to the experimental gains/losses data. Gains and losses are identified by comparing the final simulated state to initial state for each gene, e.g., to qualify as a gain, a site must have status *U* in the initial state and status *M* in the final state (as is the case for the experimental gain/loss analysis).

The total distribution for the number of gains as a function of base pair distance to the nearest *M*-site (in the same methylatable-gene-region) is compiled from all simulated genes and all replicates, along with the equivalent distribution for the number of losses. Gain (or loss) rates are calculated by dividing the distributions for number of gains (or losses) by the normalization: the distribution of the number of *U*-sites (or *M*-sites) sites in the initial state, again as a function of distance to the nearest *M*-site. Finally, we convert to an average gain and loss rate per cell cycle, as described above for the experimental case. Also, see above for further details of how we relate the modelled gain and loss rates per cell cycle to those measured experimentally. Both the modelled and experimental gain and loss rate distributions are plotted after smoothing over a 10 base pair window. In Fig. 3E,F, S6D,E,G, S10A-C, S13C ‘All experimental data sets.’ includes 8 lines from the Becker et al. dataset, 4 lines from the Schmitz et al. dataset and the *WT* dataset presented in this work.

Initially we manually fit the model parameters using only the gain/loss rate distribution to the nearest *M*-site, calculated over whole gene-body methylated genes. This dataset provided no evidence of the need to vary the strength of the cooperative interaction amplitude as a function of mean CG site density. For this fit, we therefore set *α*_*m*_ = *β*_*m*_ = 0. The fitted parameter values are given in Table S7A.

The parameter values were then revised to simulate our annotation of methylatable regions (as opposed to simulating whole genes as previously). The methylatable regions were identified using the frequency of methylation across *Arabidopsis* accessions and the Col-0 H2A.Z thresholds (as described earlier). The parameters for this model are given in Table S7B. Switching to the new annotation resulted in a ∼10-fold increase in the spontaneous *de novo* rate (to 5×10^-6^ per site per cell cycle), compared to that found using the whole-gene annotation (5×10^-7^ per site per cell cycle, Table S7A). This new spontaneous *de novo* rate is similar to the gain rate observed in UM segments of gbM genes in *WT* (3×10^-6^ per site per cell cycle for the Schmitz dataset, Table S1). The spontaneous *de novo* rate found using the whole-gene annotation is instead comparable to the gain rate observed in *WT* UM genes (6.4×10^-7^ per site per cell cycle for the Schmitz dataset, Table S1). Spontaneous *de novo* rates were fit to the tail of the gain-rate distributions (at a distance of *x* = 1000 bp, Fig. S6G and Fig. S10C).

The fit to methylation level histograms as a function of mean CG site density was, however, not satisfactory (Fig. S9C), motivating the introduction of a varying cooperative interaction amplitude as a function of mean CG site density. The final model parameterization is given in Table S7C, and was obtained by fitting to the gain/loss rates over methylatable-regions in conjunction with the simulated steady-state methylation level distribution as described below. After completing this fit, the model was tested by, for example, comparing the simulated and experimental gain/loss rate distributions as a function to the nearest *U*-site (calculated analogously to the distance to the nearest *M*-site distributions described above) (Fig. S13C). We did not fit to these distributions. Further comparisons between modelled and experimental methylation properties are described in the next section.

To investigate the relative importance of the primary and secondary components of the cooperative gain interaction, we simulated the number of gains predicted by the model over 30 generations using the parameterization in Table S7A, but with the secondary cooperative gain component 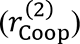 set to zero (Fig. S6D). Similarly, the strength of the cooperative maintenance contribution was assessed by simulating the number of losses predicted by the model (parameterized according to Table S7A), but in the absence of cooperative maintenance (*β*_0_ = 0) (Fig. S6E). This provides a uniform loss rate at the level of the background loss rate (*ε*). For the final model parameterization (Table S7C), we also assessed the impact of the various cooperative gain/loss interactions, again assessed over 30 generations (Table S9).

### Simulated steady-state methylation patterns

The steady-state methylation patterns predicted by the model are investigated by simulating for 100,000 generations for the specified number of replicates. In many cases, our simulations use a fully unmethylated initial state, however, we also confirmed that the model has a unique steady-state by also simulating from an experimental Col-0 or fully methylated initial state (Fig. 4B). We confirmed that a simulation time of 100,000 generations is ample for the model (parameterized according to Table S7C) to reach steady-state (Fig. S11D). As the steady-state is independent of the initial state choice (Fig. 4B), when using the experimental Col-0 initial state we arbitrarily assign an initial state of *U* to the CG sites with unknown methylation status in the Col-0 consensus state. We compare the modelled steady-state methylation states, for a certain number of replicates, to those observed for an equivalent number of *A. thaliana* accessions, using the 1001methylomes resource (Kawakatsu et al., 2016). Each independently simulated replicate is taken to represent a single accession, to investigate the extent to which our stochastic model of methylation dynamics can account for the observed natural variation in gene body methylation level across Col-like accessions (defined above). All Col-like accessions are assumed to have genetically equivalent methylation machinery. This approach neglects all effects of selection pressure that may affect the observed methylation patterns and assumes that no significant changes occur to the methylation machinery, or H2A.Z distribution, over the timescale required for gene-body methylation to reach steady-state. We also neglect any possible outcrossing between different accessions over this timescale: such outcrossing may occasionally occur, but provided it is between the same accession or with another Col-like accession, then statistically equivalent methylomes will be combined, which will not change the steady-state methylation distributions.

Additionally, we investigate the sensitivity of our model to potential genetic diversity in the methylation machinery, by adjusting the overall strength of the cooperative interaction pathway (parameterization details discussed above) and/or spontaneous *de novo* strength. Simulated methylation patterns are compared to *A. thaliana* methylation-outlier accessions (see above), assuming the same methylatable-regions as identified for Col-like accessions (Fig. 6A-D, S15). We note, however, that methylation-outlier accessions may also have accession-specific methylatable-regions.

For each methylation-state (either a simulated replicate, or measured *A. thaliana* accession), we calculate the mean methylation level, ⟨*M*〉, for each methylatable-region, where 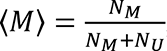, with *N_M_* and *N_U_* being the number of CG-sites identified with status *M* and *U* respectively in that methylatable region. For the simulated states, this calculation is performed at the end of the simulated period (100,000 generations). The total number of CG-sites in the simulations of the methylatable region is *N*_*CG*_ = *N*_*M*_ + *N*_*U*_, by definition. This is not the case for the observed methylation states, where some CG-sites are assigned a status *X*, either due to having an indeterminate methylation status (as described previously), or a SNP occurring at that site, so that *N*_*CG*_ = *N*_*M*_ + *N*_*X*_ + *N*_*U*_. In the rare event that *N*_*M*_ + *N*_*U*_ = 0 for a methylatable region in an accession, we assign ⟨*M*⟩ = 0. Histograms of the distribution of ⟨*M*⟩ (Fig. 4A-B,6A-D, Fig. S7, S9B-C, S10E-H, S12, S15) are normalized to the number of replicates (either simulated or observed accessions). The clear spike at ⟨*M*⟩ = 0 corresponds to fully unmethylated regions. In addition, extra structure in this distribution arises due to the fractional definition of methylation level: a value of ⟨*M*⟩ = 1/2 can be obtained for any gene-region with an even value of *N*_*CG*_, whereas a value of approximately ⟨*M*⟩ = 49/100, for example, can only be obtained from a much more limited subset of gene-regions. To study the variability of methylation levels between replicates, we also calculated the standard deviation of ⟨*M*⟩, *σ*_⟨*M*〉_, over all replicates for every gene-region.

We manually fit the final model-parameterization (Table S7C), which includes a linear variation of the cooperative interaction strength as a function of mean CG-site density across the gene-region, simultaneously to both the gain and loss rates measured over 30 generations (as described previously) (Fig. S10A-C) and the distribution of ⟨*M*⟩ at steady-state (Fig. 4A, S9C). To facilitate this fit, distributions of ⟨*M*〉 (not shown) were examined in greater detail for subsets of gene-regions, grouped according to both the length of the gene-region, *L*_*locus*_, and the mean CG-density, *ρ*_*CG*_, as defined previously. Subgroups were defined as: *L*_*min*_ ≤ *L*_*locus*_ < *L*_*max*_ for *L*_*min*_ = {0, 1 × 10^3^, 2 × 10^3^, 4 × 10^3^, 6 × 10^3^, 1 × 10^4^} bp and *L*_*max*_ = {1 × 10^3^, 2 × 10^3^, 4 × 10^3^, 6 × 10^3^, 1 × 10^4^, 3 × 10^4^} bp and 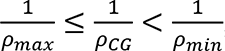, for 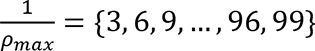 bp and 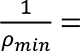 {6, 9, 12, …, 99, 102} bp, with *ρ*^−1^ increasing in units of 3 bp. We compared the simulated steady-state methylation distributions to 30 Col-like replicates, selected for their high sequencing coverage (described above).

Adjusting the value of the spontaneous background rate (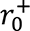) was instrumental in capturing the height of the spikes at the origin. It was challenging to accurately capture the increase in mean methylation level with increasing locus length (⟨*M*⟩[*L*_*locus*_]). We found that slight adjustments to the power-laws (∼ ± 5%) were possible while still maintaining a good fit to the gain/loss rate distributions, while having a noticeable effect on ⟨*M*⟩[*L*_*locus*_]. The other influential parameters were: *ε*, *α*_*c*_ and *α*_1_, though the latter was well constrained by the gain/loss rate distributions (as were the plateau lengths and power-law divergences), while the sensitivity to *ε* was similar to that of the power-laws. After fixing all other parameters, an initial estimate of *α*_*m*_, *α*_*c*_, *β*_*m*_ and *β*_*c*_ was found by independently manually fitting the individual sets of histograms generated with a fixed value of *ρ*_*CG*_ to a value of *α*_0_ and *β*_0_ before then using a linear fit to the resulting values of *α*_0_(*ρ*_*CG*_) and *β*_0_(*ρ*_*CG*_) to extract corresponding values of *α*_*m*_, *α*_*c*_, *β*_*m*_ and *β*_*c*_.

The fully parameterized model was then simulated to steady-state (100,000 generations) for 740 replicates and compared to all non-redundant Col-like accessions with sequencing coverage > 50% (see above). The performance of the model at the individual gene level was tested by comparing the predicted and observed values of both ⟨*M*⟩ and *σ*_⟨*M*⟩_, both averaged over the 740 simulated/observed accession replicates respectively, for each individual methylatable region (Fig. 4D, 5D, S12).

We also tested the model’s ability to produce spatially similar methylation patterns to those observed, by calculating the distribution of the following neighbor-pair separations in units of base pairs: {*MM*, *MU*, *UU*, *XX*} neighboring sites. Here, an *XX*-neighbour-pair is defined to be a pair of neighbouring CG sites, at least one of which cannot be identified as either *M* or *U* from the sequencing data. Note that there are no *XX*-neighbour pairs, by definition, for the simulated states. All three simulated distributions are then normalized to the total number of pairs (of any type), totaled over all 740 realizations. Equivalently, the four experimental distributions are all normalized to the total number of pairs (of any type), totaled over all 740 accessions. The three simulated distributions are shown directly (Fig. S13D). For the experimental distributions, we add the *XX*-neighbour pairs distribution to each of the other three to produce the green bands. The bottom of each green band therefore corresponds to the case that none of the *XX*-neighbour pairs (if their methylation status were known) contribute to the depicted distribution. The top of each green band corresponds to the case that all the *XX*-neighbour pairs contribute to the depicted distribution. The simulated distributions are therefore expected to lie somewhere within the green bands. Although each *XX*-neighbour pair can only actually belong to one of the three distributions, the fraction of *XX*-neighbour pairs that corresponds to each distribution will vary considerably as a function of pair separation. This is because the sites with unknown methylation status are not evenly distributed, but instead are concentrated close to existing methylation.

### Assessing the scale of simulated methylation fluctuations

The fluctuations in methylation level were investigated by simulating 740 replicates, starting from the all-*U* initial state each time. The time of the first methylation gain was recorded for each locus, and averaged over the 740 replicates. This is compared to the theoretically expected value of: 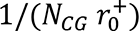 cell cycles, or 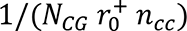 generations (red curve in Fig. S11B). A steady-state methylation level was found for each replicate of each locus, by first simulating (from the all-*U* initial state) for an equilibration-time of 50,000 generations (ample time to reach steady-state, Fig. S11D). The simulation was then continued for another 50,000 generations (denoted by *T* below), over which the overall methylation level of the locus, ⟨*M*⟩, was time-averaged to find the steady-state methylation level, 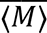, where:

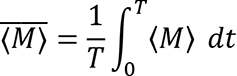

The time for an individual locus to first reach steady-state is then defined as the time the methylation level of that locus first reached the value of 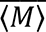. This calculation is repeated independently for each replicate of each locus, with the mean and standard deviation over replicates then shown in Fig. 4C and Fig. S11C respectively. An estimate of the typical magnitude of fluctuations away from the mean steady-state methylation level is found from the standard deviation of the methylation level over time, Σ, where:

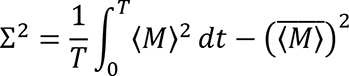

Again, this quantity is calculated individually for each replicate of each locus, with the mean over replicates shown in Fig. S14C. Finally, we study the ‘greatest’ fluctuation occurring in the second 50,000 generations of the above 100,000 generation simulations. Here we use ‘greatest’ to refer to the fluctuation showing the largest deviation in methylation level from the time-averaged mean. Both the magnitude and duration of this fluctuation are recorded for each replicate of each locus, with the mean and standard deviation over replicates shown in Fig. 5B and Fig. S14A respectively for the magnitude, and in Fig. 5C and Fig. S14B respectively for the duration. We note that the magnitude of the greatest fluctuation will depend on the length of the time-interval, *T*, that is studied: the longer the time-interval, the higher the likelihood that more extreme fluctuations will be observed.

### Fits to mutant mean gain and loss rates over methylatable regions

The large fluctuations and insufficient range, particularly in the observed mutant loss rate distribution (as a function of distance to the nearest *M*-site), preclude a full fitting to the gain/loss rate distributions (Fig. S5D and Fig. S8C). Instead, we therefore used the observed mean gain/loss rate over methylatable regions (Table S1). Although the simulated mean gain/loss rates for the full model fit to Col-0 are close to those found for our *WT* data set (Fig. S10D; Table S1;S9), there are still ∼10% discrepancies. We therefore calculated a target mean gain and loss rate for each mutant (Table S10) by applying the same relative change from the simulated mean *WT* rates as is found from comparing the mean rate for each mutant to the corresponding newly measured mean *WT* rates (as these were all measured using a consistent experimental design). The simulated gain/loss rate distributions were found by fitting the overall cooperativity strength (*r*^∗^), and the background loss rate (*ε*), to the target mean gain/loss rates.

As relatively recently generated mutants are used, we assume that their methylation state is still very close to that of the Col-0 consensus state. Consequently, as before, we use the Col-0 consensus state for the initial state and simulate for 30 generations, using 100 replicates to find the simulated mean gain/loss rates over methylated regions. Parameters are as for the full model (Table S7C) but with the overall cooperativity strength (*r*^∗^) and background loss rate (*ε*) adjusted according to Table S10. For *h2az,* caps were introduced to the cooperative interaction to ensure that 0 < 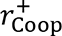 < 1 and: 0 < *γ* < 1. Using these parameter values, the steady-state methylation level distribution, and the time to reach steady-state were then found for each mutant, similarly to the description above for Col-like accessions, though only using 30 modelled replicates for each simulation and from the Col-0 consensus initial state.

Finally, we note that for the *ros1* mutant, we could alternatively produce a very similar fit to that produced by altering overall cooperativity strength by adjusting the spontaneous *de novo* rate (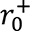) instead of the overall cooperativity strength. These results are not shown, as they did not appreciably alter the steady-state state methylation level. We emphasize that all fits to mutant rates are tentative due to the restricted amount of data available for the fit (only a single average gain and loss rate per mutant, as opposed to the spatially resolved Col-0 gain/loss rate distributions and the steady-state methylation levels) available for the *WT* population. Long-timescale (steady-state) simulations only reflect the direct consequences of the altered short-timescale (30 generation) methylation dynamics. Time to steady-state calculations are estimates, as it is unknown which side of the steady-state each locus is in the initial state and therefore loci are assumed to transition in the direction of the overall mean of the distribution. Any potential indirect effects, such as altered methyltransferase targeting are not accounted for.

### Use of previously published datasets

GSE39045: H2A.Z ChIP-seq data from (Coleman-Derr & Zilberman, 2012)

SRA035939: 30 generation epiRIL BS-seq data from (Schmitz et al., 2011)

PRJEB2678: 30 generation epiRIL BS-seq data from (Becker et al., 2011)

GSE178684: Generational bs-seq data from (Hazarika et al., 2022)

GSE64463: Generational bs-seq data from (Shahryary et al., 2020)

GSE43857: 1001 Epigenomes bs-seq data from (Kawakatsu et al., 2016)

GSE96994: Nucleosome positions from (Lyons & Zilberman, 2017)

## Author Contributions

A.B. developed the mathematical model, performed analysis, and wrote the paper E.H. performed DNA methylation experiments and analysis, and wrote the paper Z.S. performed population genetics analyses and wrote the paper J.D.M. performed DNA methylation and population genetics analyses, and wrote the paper D.B.L. performed nucleosome positioning and DNA methylation analyses M.H. conceived the research plan, supervised analysis, and wrote the paper D.Z. conceived the research plan, supervised experiments and analysis, and wrote the paper

## Acknowledgements

We would like to thank Xiaoqi Feng, Ander Movilla Miangolarra and Suzanne de Bruijn for discussions. This work was supported by BBSRC Institute Strategic Programme GEN (BB/P013511/1) to M.H. and D.Z, and by a European Research Council grant MaintainMeth (725746) to D.Z.

## Supplemental Figure Legends

**Supplemental Figure 1.**
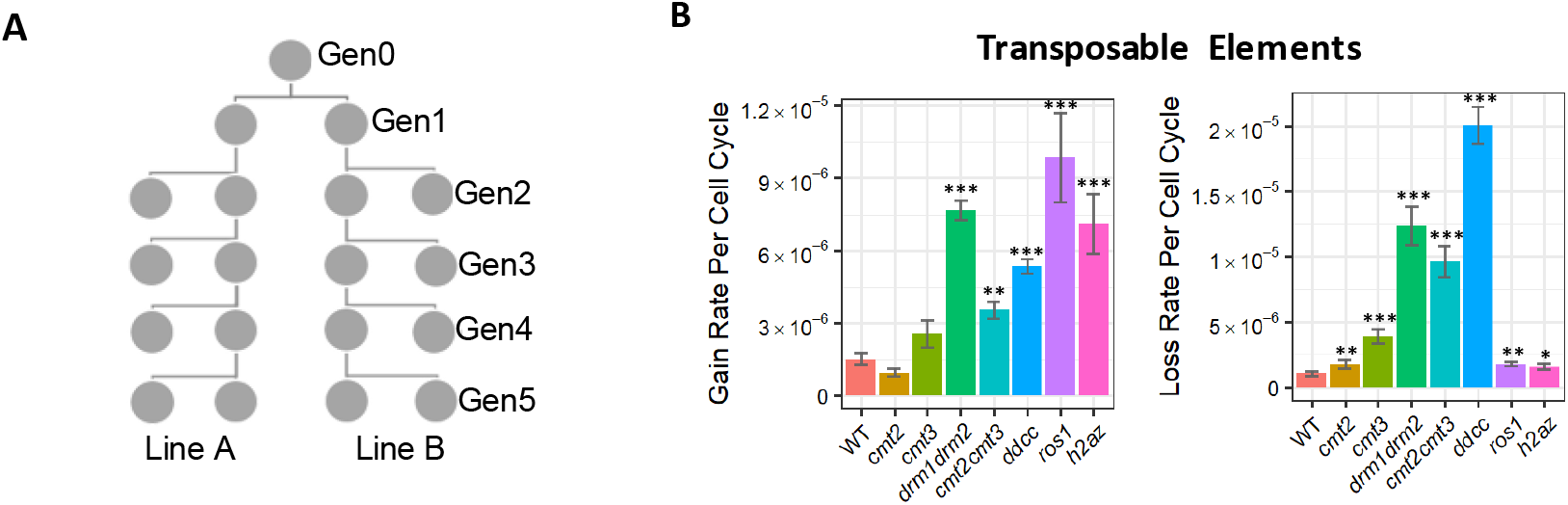
**A)** Pedigrees of *WT* plants used. Two offspring of an individual parent plant of each genotype were sampled at each generation for between four and six consecutive generations. **B)** Per cell cycle rates of mCG gain and loss at individual CG sites within transposable elements. Loss rates increase significantly in all mutants. Gain rates cannot be measured accurately due to the low proportion of uCG sites. A significant difference from the *WT* rate is indicated by *p=0.01 to 0.001, **p=0.001 to 0.00001, and ***p<0.00001 (Fisher’s exact test). Error bars indicate standard error.

**Supplemental Figure 2.**
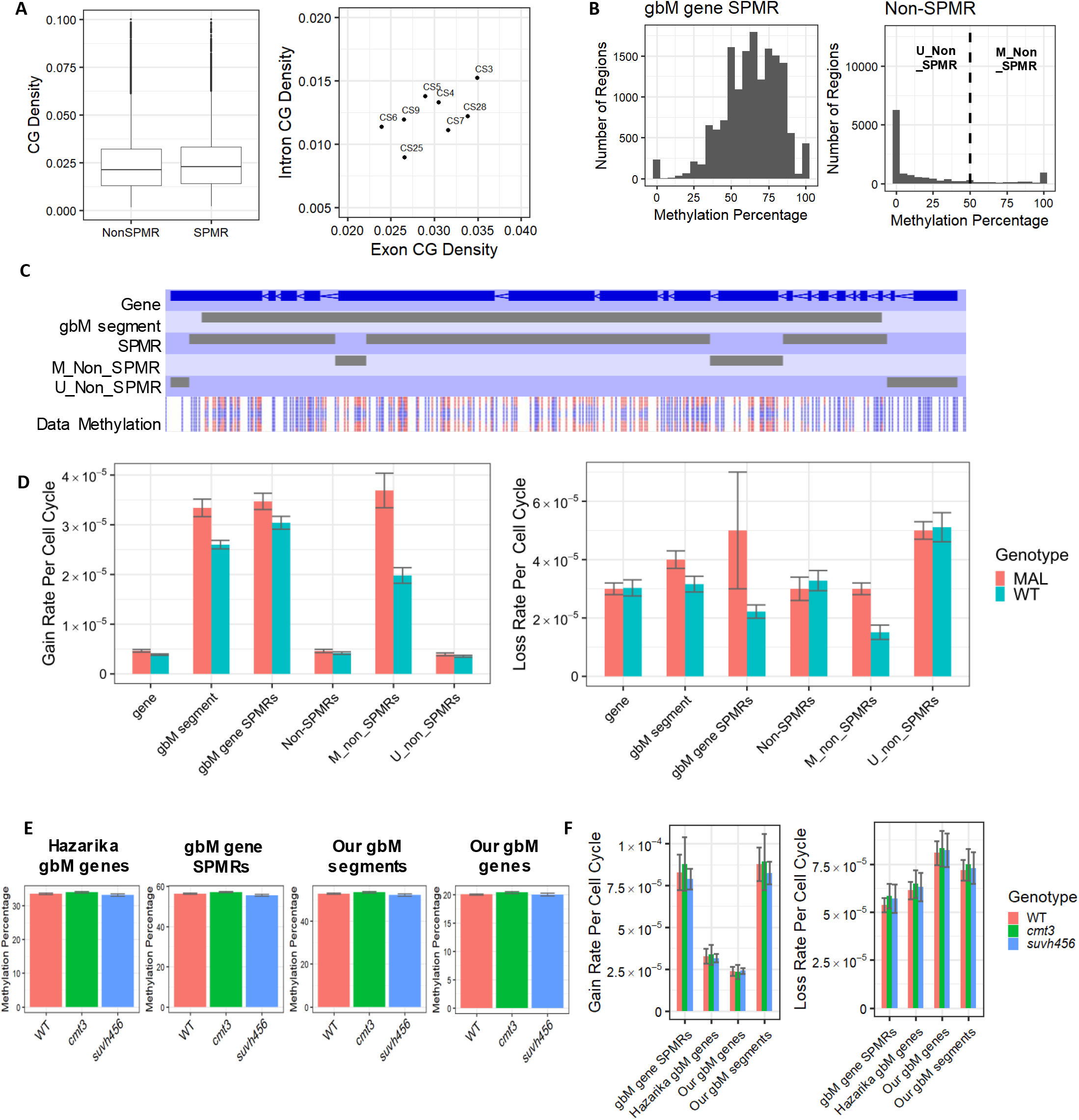
**A)** Reanalysis of SPMR CG site density demonstrates that it is similar to that of non-SPMRs (left), with introns having a lower density than exons (right), and generally much lower than previously reported (Hazarika et al., 2022). This is true for all SPMR chromatin states (CS) described in (Hazarika et al., 2022) (right). **B)** Regions of gbM genes outside of SPMRs (non-SPMRs) are bimodal with respect to mCG and fall into two groups: M_non_SPMRs (those with mCG ≥ 50%) and U_non_SPMRs (those with mCG < 50%). **C)** M_non_SPMRs and SPMRs largely fall within our gbM segments. A genome browser display of AT1G03060 is an example of this. Red indicates methylated reads and blue indicates unmethylated reads. **D)** Epimutation rates – especially gain rates – in methylated regions (gbM segments, SPMRs and M_non_SPMRs) are similar to each other but distinct to unmethylated regions (U_non_SPMRs). Rates calculated using published data and 30 generation analyses (MAL) are similar to our new data and individual generation analyses (WT). Non-SPMRs are composed of two groups (M_non_SPMRs and U_non_SPMRs) but, as U_non_SPMRs contain 9-fold more CG sites than M_non_SPMRs, gain rates in non-SPMRs are dominated by the unmethylated U_non_SPMRs, and thus the rates are similar between U_non_SPMRs and all non-SPMRs. Error bars indicate standard error. **E)** Methylation levels in *WT*, *cmt3* or *suvh4/5/6* samples from (Hazarika et al., 2022) are effectively indistinguishable. Methylation level calculated per sample in each region by calculating the number of C reads divided by the number of total reads as a percentage at each site, and averaging this across the region. Plot shows mean methylation level averaged across all samples of a given genotype, with error bars representing standard error. C reads represent methylated reads of a given cytosine in bisulfite sequencing experiments. No significant differences were observed between genotypes (p<0.01, Student’s t-test). **F)** Rates of methylation gain and loss in *WT*, *cmt3* or *suvh4/5/6* samples in gbM genes and SPMRs from (Hazarika et al., 2022), in our gbM segments and in our gbM genes, all using Hazarika et al. sequencing data. Some samples were of insufficient coverage (<10X) to make methylation calls (without use of imputation, which determines methylation status based on the methylation status of nearby cytosines and is unsuitable for this analysis) and thus these lines were excluded. Error bars indicate standard error. No significant difference from the *WT* rate is observed in gbM gene SPMRs, published gbM genes (Hazarika gbM genes), gbM segments or our gbM genes for *cmt3* or *suvh456* mutants (p<0.01, Fisher’s exact test).

**Supplemental Figure 3.**
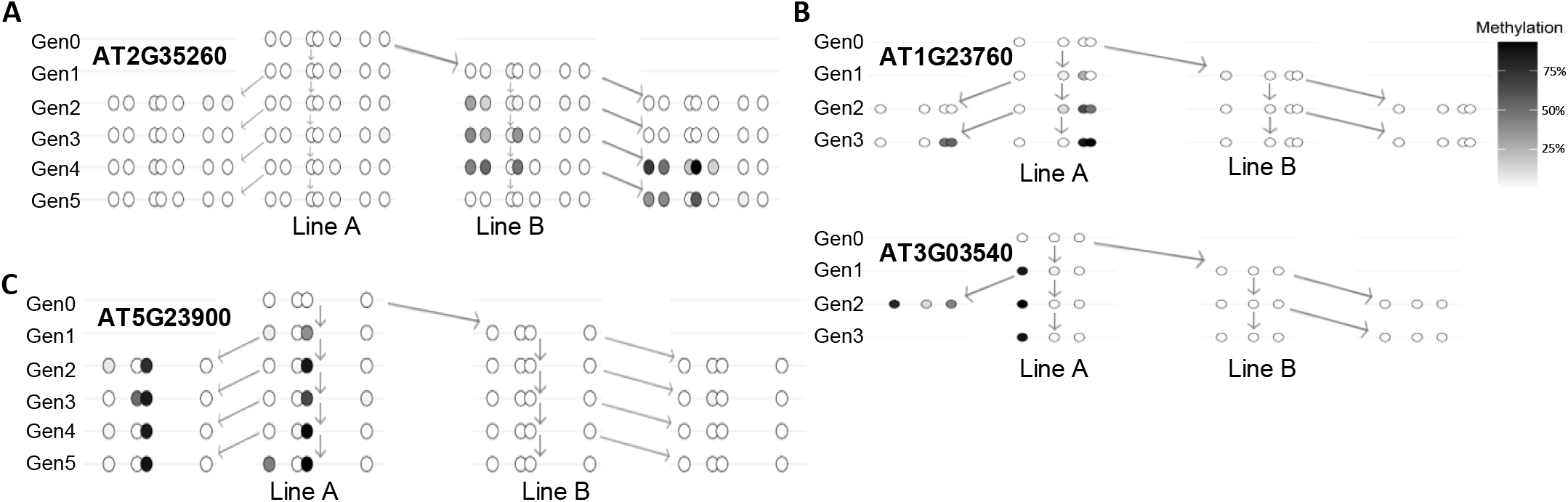
**A-C)** New methylated clusters occur in previously entirely unmethylated genes in *drm1drm2* (**A**), *ddcc* (**B**), and *cmt3* (**C**). Each circle represents an individual CG pair, with darkness of fill indicating methylation percentage.

**Supplemental Figure 4.**
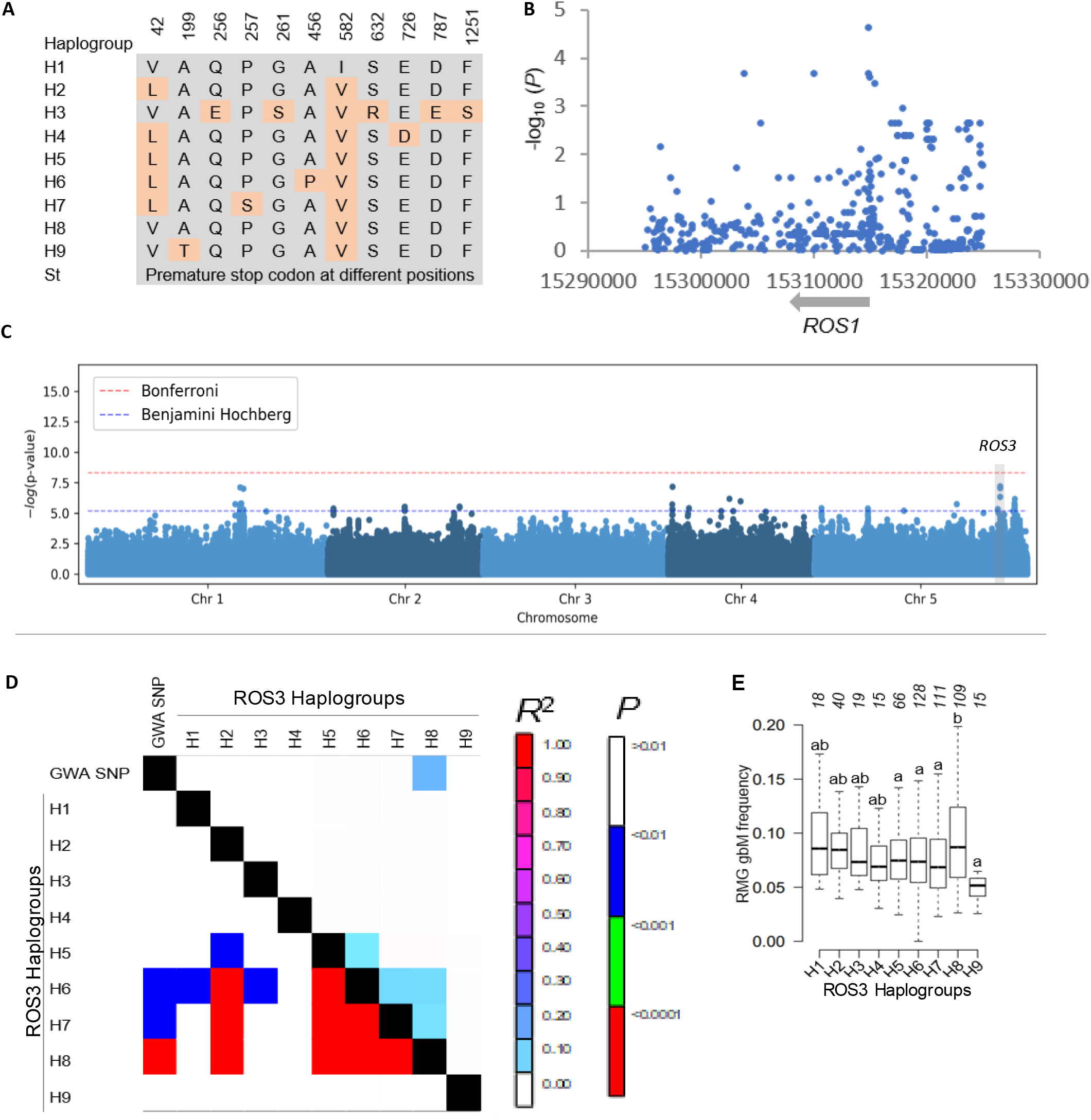
**A)** Position and nature of amino acid polymorphisms among *ROS1* haplogroups. **B)** Manhattan plot for association P value of SNPs with gbM frequency of RMGs in *ROS1* region. Each dot corresponds to a SNP. **C)** Manhattan plot of genome-wide association mapping for gbM frequency of RMGs in *Arabidopsis* accessions. Horizontal dashed blue and red lines respectively show the 0.05 False Discovery Rate and Bonferroni thresholds. **D)** Linkage disequilibrium (R^2^) analysis revealed that ROS3 haplogroup 8 segregates together with GWA SNP Chr5:23536319 in *Arabidopsis* populations. The color of different cells of the heatmap denotes the value of R^2^ and P values as indicated in the color key. Linkage disequilibrium between ROS3 haplogroups and GWA SNP was calculated using TASSEL. **E)** GbM frequency of RMGs in haplogroups of ROS3. Amino acid sequences were aligned and accessions invariant for amino acids were classified into a haplogroup. Different letters indicate significant differences at p < 0.05 (one-way ANOVA), i.e. groups denoted with “a” are statistically different from “b” and ab ones are different from neither a nor b. The number of accessions is indicated for each haplogroup (top).

**Supplemental Figure 5.**
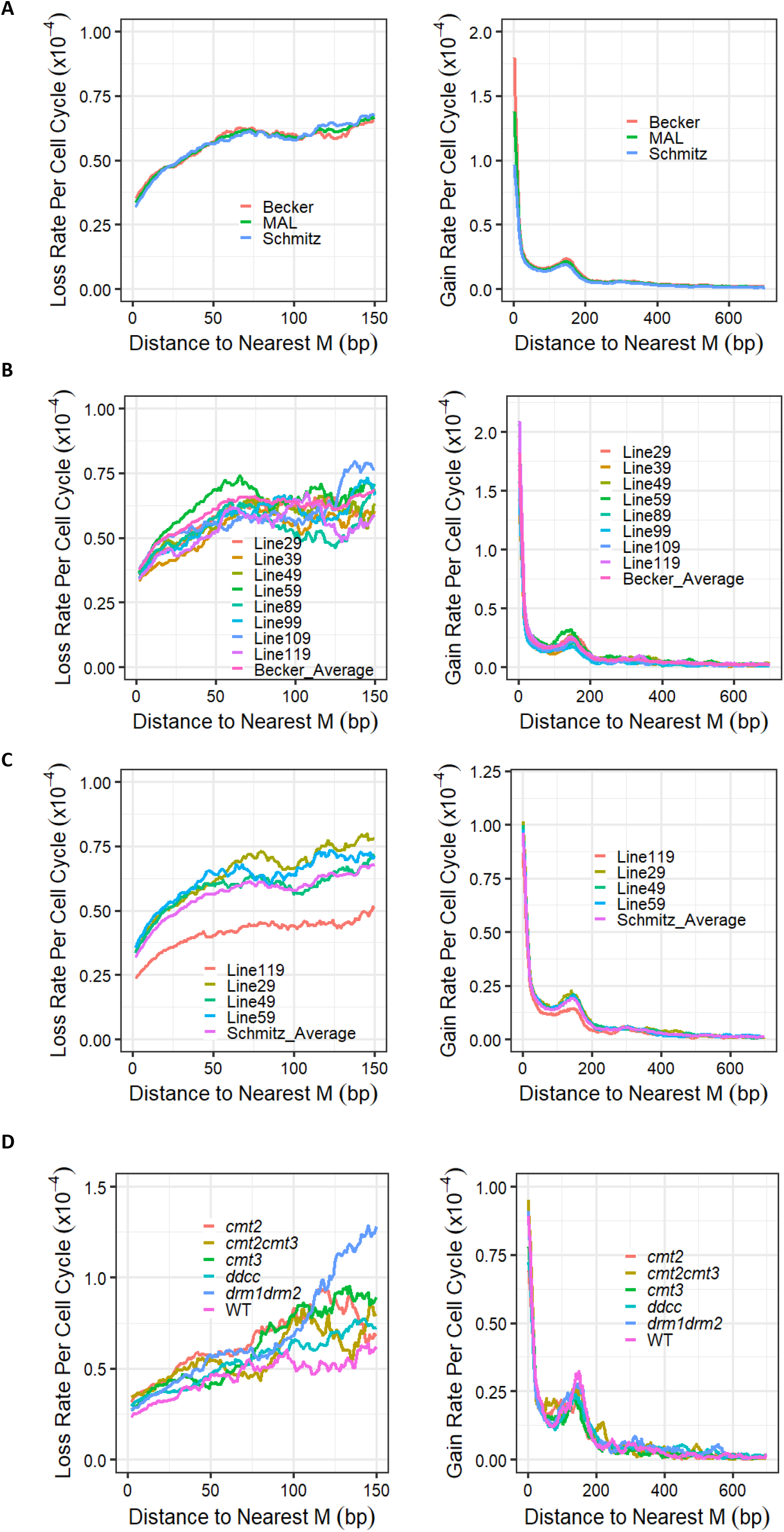
**A)** Profiles of methylation gain/loss rates relative to the nearest mCG in gbM genes are similar in the different datasets of previously published mutation accumulation line (MAL) data (Becker et al., 2011; Schmitz et al., 2011). Rates plotted over whole gbM genes and shown as a 30 bp moving average. **B-C)** Different lines within the Becker et al., 2011 (**B**) and Schmitz et al., 2011 (**C**) datasets show similar profiles to each other for both gains and losses, plotted relative to the nearest mCG over whole gbM genes. Rates shown as a 30 bp moving average. **D)** Profiles of methylation gain/loss rates relative to the nearest mCG remain similar in mutants of all non-MET1 methyltransferases. Rates plotted over whole gbM genes and shown as a 30 bp moving average.

**Supplemental Figure 6.**
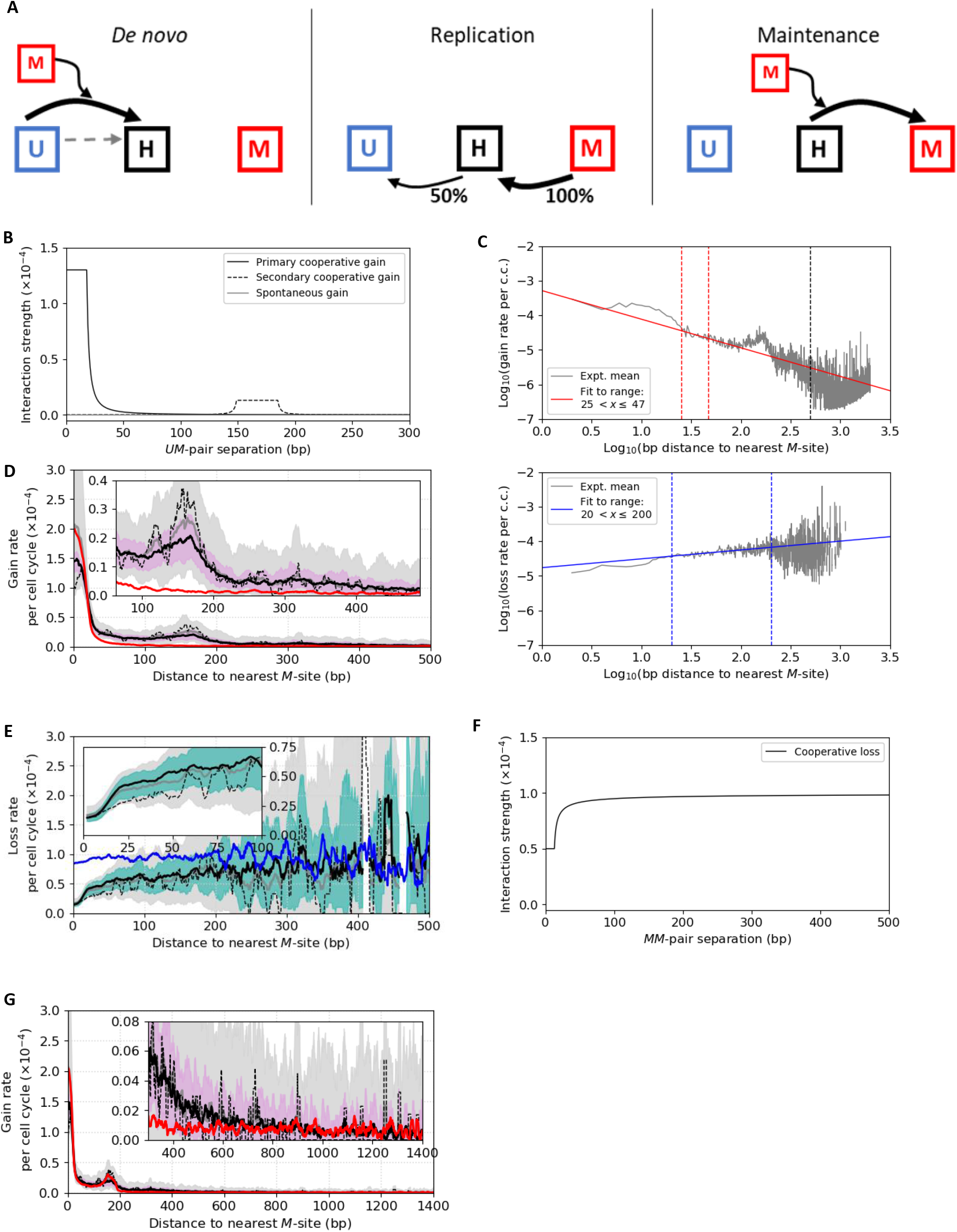
**A)** Underlying three state model (detailed description in Methods). **B)** Gain-interaction components of the effective two state model. Cooperative gain strength is modelled using a power-law decay as a function of base-pair distance between each *UM*-pair of CG sites. The primary interaction component (solid black) is strongest for proximate CG sites, whereas the secondary component (dashed) peaks for CG sites separated by ∼170 bp reflecting the enhanced likelihood of gains occurring at uCG sites that are approximately one nucleosome spacing distance from existing mCG sites. The total cooperative interaction is a sum of contributions from all *UM*-pairs containing the target uCG site, within a single locus. A uniform background gain rate independent of existing methylation (grey) is also included, shown here increased by a factor of 100 for clarity. **C)** Col-0 gain (top) and loss (bottom) rates (for whole gbM genes, mean of four replicates from (Schmitz et al., 2011)) plotted as a function of distance to the nearest mCG site using a log-log scale (grey lines). Colored straight lines are linear fits (signature of power-law behavior) to data within the regions indicated by vertical colored dotted lines (gains (red): 25 < *x* < 47 bp, losses (blue): 20 < *x* < 200 bp). Loss rates are consistent with a power-law form over at least one decade. Gains rates are more subtle due to the presence of two interaction components. The choice of a narrow fitting window isolates the gain rate data dominated by the primary interaction component. Extrapolating the corresponding fit to long length-scales (beyond the secondary interaction peak around 170 bp) is consistent with the observed gain-rate at around *x* = 500 bp (dotted black line). **D)** Gain rate profile as a function of distance to the nearest mCG site (calculated over whole gbM genes). Simulated and experimental data plotted as in Fig. 3F, except here simulation output (red) only includes the primary cooperative interaction component: this is insufficient to reproduce the enhanced gain rate at ∼170 bp (highlighted in inset). **E)** Loss rate profile as a function of distance to the nearest mCG site (calculated over whole gbM genes). Simulated and experimental data plotted as in Fig. 3E, except here simulation output (blue) includes no cooperative component to the maintenance interaction, resulting in a ∼5-fold increase in losses at short distances. Inset highlights the discrepancy between simulation and data at these short length-scales. **F)** Cooperative loss interaction strength as a function of CG-site separation for an individual *MM*-pair. At long length-scales there is a power-law convergence to a constant interaction strength. The total cooperative loss rate at a given CG site is formed from the product of these individual loss probabilities multiplied by the overall scale of the effective maintenance interaction (10^-4^ per cell cycle for the fit to whole gbM genes). Detailed derivation for cooperative maintenance interaction given in Methods. **G)** Data and simulation for gains (as in Fig. 3F) fit well at around 1000 bp, determining the magnitude of the uniform background gain rate.

**Supplemental Figure 7.**
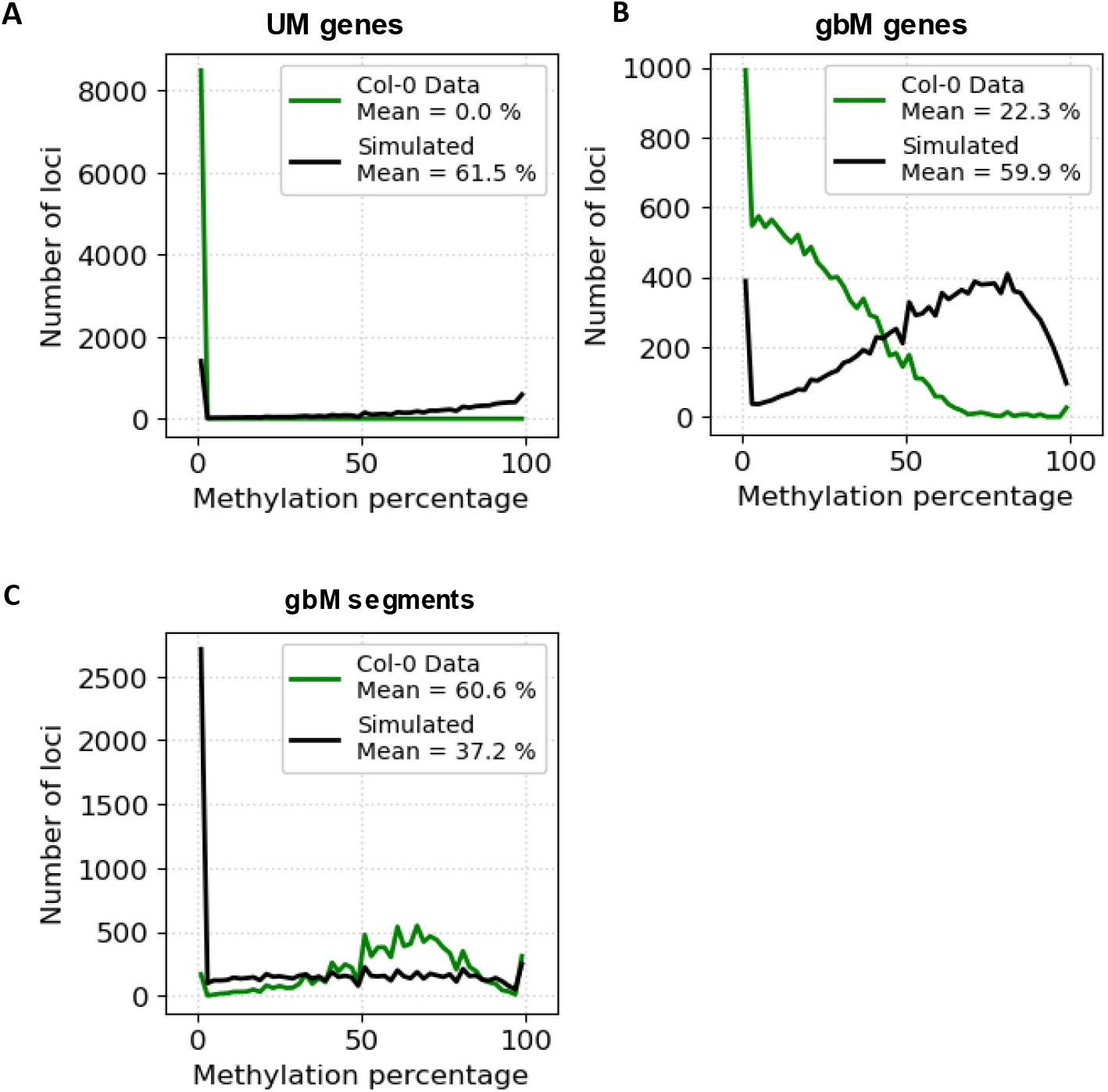
Distribution of methylation levels for unmethylated genes **(A)**, gbM genes **(B)**, and gbM segments **(C)** within Col-0 (green) and for the corresponding simulated steady-state (black). Whole gbM genes and UM genes both exhibit over-methylation in simulated steady-state, whereas gbM segments exhibit severe under-methylation in simulated steady-state. Average methylation for each dataset is shown. Simulations run for 100,000 generations from Col-0 initial state and normalized for 30 realizations (model parameters given in Table S7A).

**Supplemental Figure 8.**
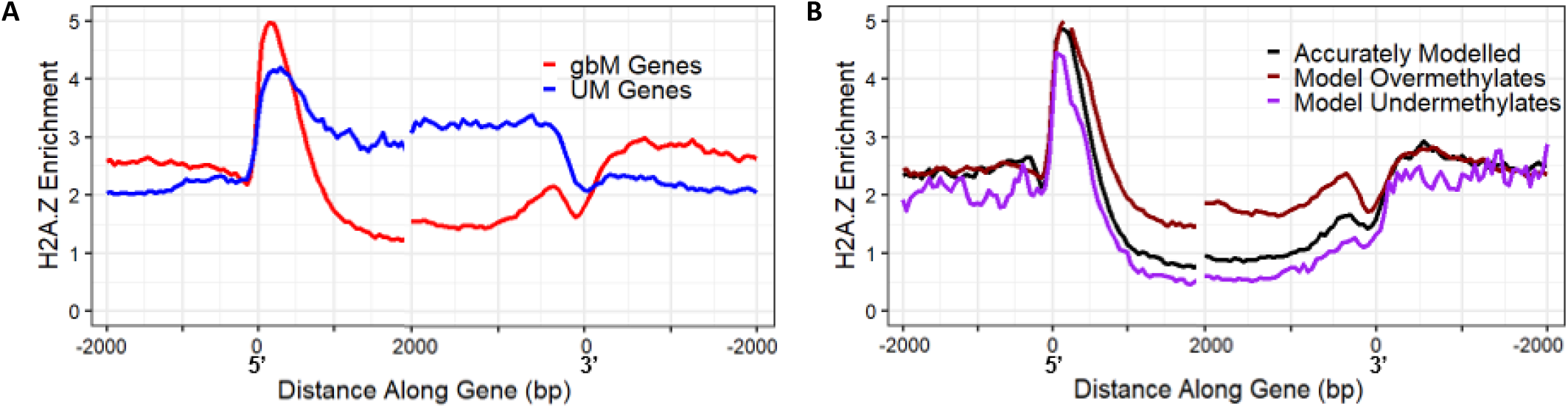
**A)** H2A.Z enrichment anticorrelates with methylation levels, higher in unmethylated (UM) genes than gbM genes and higher at 5’ gene ends than within genes. Left panel displays H2A.Z enrichment around the 5’ end, indicated by 0; right panel displays H2A.Z enrichment around the 3’ end, indicated by 0. H2A.Z enrichment calculated as IP-input and averaged over gbM genes (red, N=14,581) or UM genes (blue, N=12,045). **B)** Genes over-methylated by the model have higher H2A.Z enrichment. Mean H2A.Z enrichment shown along three groups of loci (Model Overmethylates (modelled methylation >20% above data methylation, N=8142, brown), Model Undermethylates (modelled methylation >20% below data methylation, N=361, purple) and Accurately Modelled (<20% difference between model and data methylation, N=2852, black) shown over the 5’ (left) and 3’ (right) end of the gene, indicated by 0. Simulation as in Fig. S7.

**Supplemental Figure 9.**
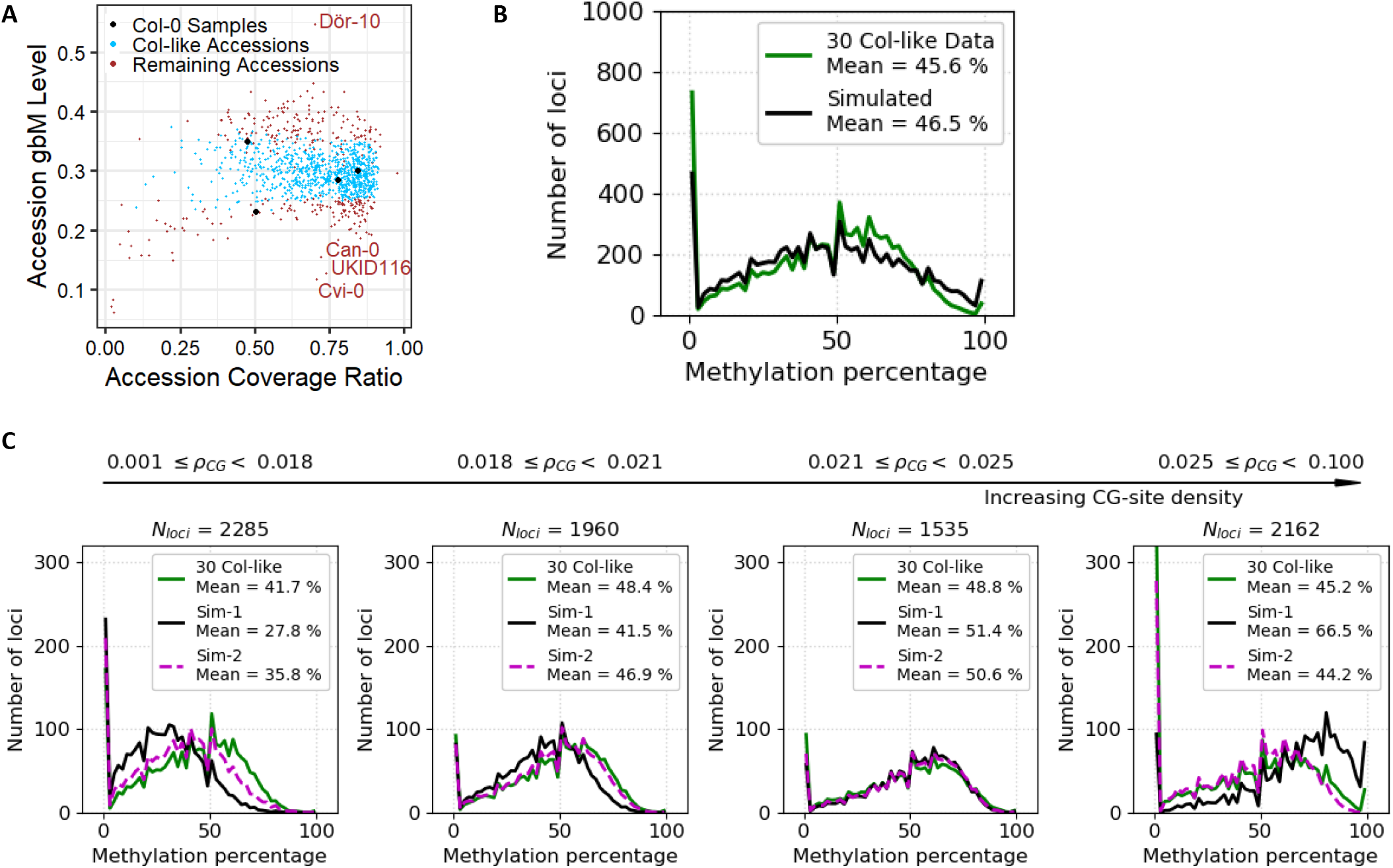
**A)** GbM levels among the accessions within the 1001 methylomes dataset (Kawakatsu et al., 2016). GbM levels in Col-0 samples (black) vary substantially. Col-like accessions were selected (blue, N=891) to be within 1 standard deviation of gbM mean. **B)** Distribution of methylation levels within high coverage Col-like accessions (N=30, green) and simulated steady-state (black) over methylatable regions. Mean for each dataset is shown. Simulations normalized for 30 locus replicates after 100,000 generations starting from Col-0 initial state (model parameters given in Table S7B). **C)** Distributions of methylation levels for loci within high coverage Col-like accessions (N=30, green) and simulated steady-state over methylatable regions either without the CG density dependent linear correction (black, model parameters given in Table S7B) or with the correction (purple dashed, model parameters given in Table S7C), split by average CG density (*ρ*_*CG*_) of locus. As density increases (left to right), loci become overmethylated by the model in the absence of the CG density correction. Mean for each dataset shown. Simulations normalized for 30 locus replicates after 100,000 generations starting from Col-0 initial state.

**Supplemental Figure 10.**
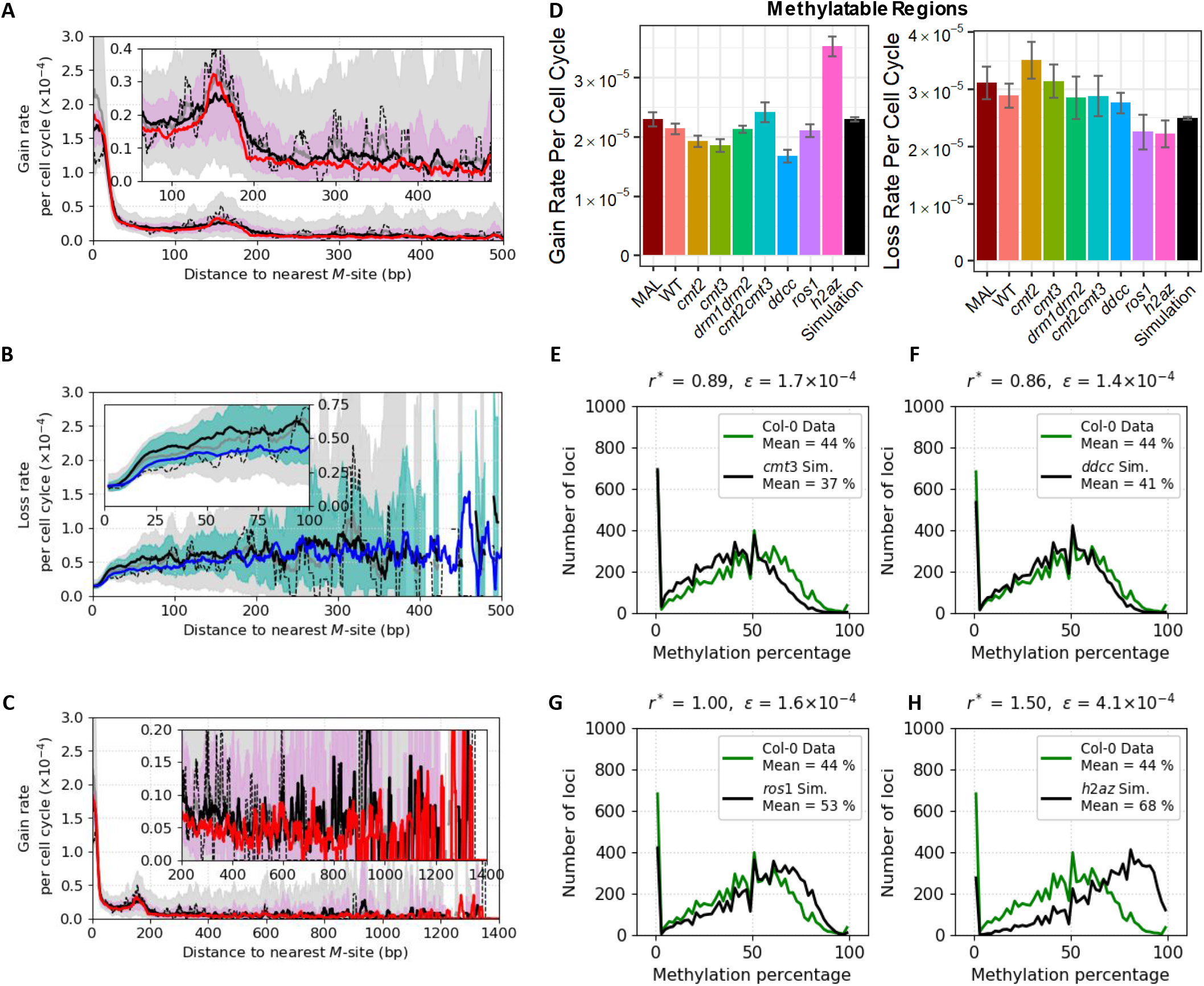
**A-C)** Profiles of methylation gain **(A, C)** and loss **(B)** rates per cell cycle, plotted as a function of distance to the nearest mCG site, calculated over methylatable regions. Datasets and simulations labelled as described in Fig. 3E,F. Rates are shown as a 10 bp moving average. Insets highlight: enhanced gain rate at a length-scale of ∼170bp **(A)**, the loss rate at short length-scales **(B)**, gain rate at long length-scales **(C)**. Simulations of full model run for 30 generations, averaged over 4 realizations, using Col-0 as initial state. Model parameters given in Table S7C. **D)** Epimutation rates in simulation over modelled methylatable regions agree well with those from experimental data. Simulation rate analysis for full model performed over 30 generations starting from Col-0 initial state using 4 simulation realizations and averaging, in order to closely resemble methodology used to calculate MAL rates. Error bars indicate standard error. Model parameters given in Table S7C. **E-H)** Distribution of methylation levels within Col-0 (green) and simulated steady-state (black) using model parameters derived from *cmt3* **(E)**, *ddcc* **(F)**, *ros1* **(G)**, and *h2az* **(H)** over methylatable regions. Model parameters used for each mutant and mean for each dataset are shown. Simulations normalized for 30 locus replicates after 100,000 generations starting from Col-0 initial state (model parameters given in Table S10).

**Supplemental Figure 11.**
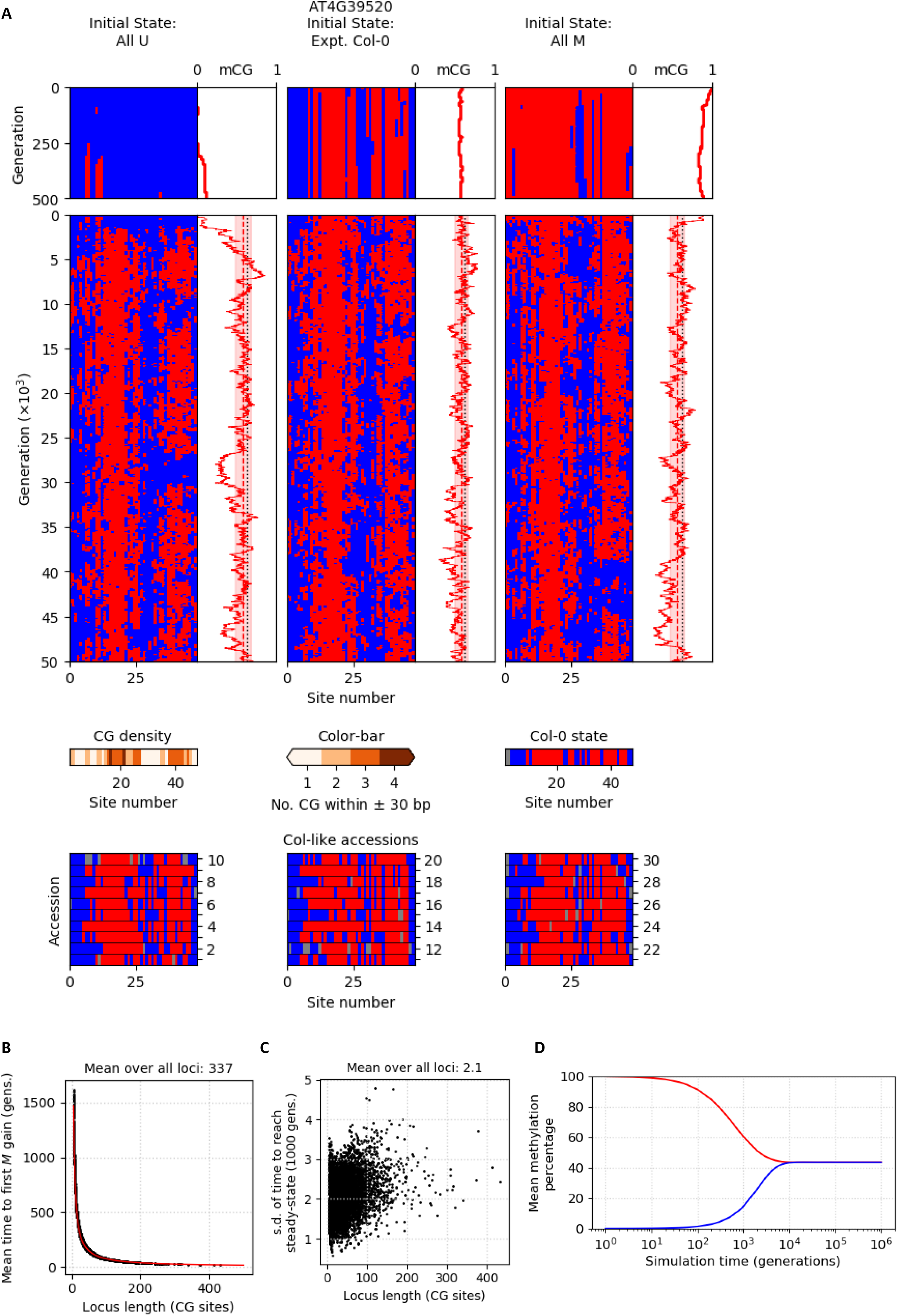
**A)** Full model simulated time-courses of an example locus (AT4G39520, 48 CG sites) reaching steady-state from either all-*U*, experimental Col-0 or all-*M* initial states. Pink highlighted windows of mCG level indicate the typical magnitude of mCG fluctuations from steady state (calculated by sd of fluctuations), and the red dashed vertical line indicates the time-averaged steady-state mCG level. Black dotted vertical line shows the average mCG level of 30 Col-like accessions. Middle: local CG density over a 60 bp window centered on each CG site. Bottom: methylation patterns for Col-0 and 30 Col-like accessions. *M*-site: red; *U*-site: blue; unknown methylation status: grey; model parameters in Table S7C. **B)** Mean time to first methylation gain from an all-*U* initial state shown for loci of different lengths, averaged over 740 locus replicates from full model simulations (model parameters in Table S7C). **C)** Standard deviation across 740 locus replicates of time to reach steady state from an all-*U* initial state for loci of different lengths from full model simulations (model parameters in Table S7C). Time shown in 1000 generations. **D)** Time to convergence of full model (in generations). Model is converged once simulations from the all-*M* initial state (red) and the all-*U* initial state (blue) reach the same steady-state. At each time point, methylation level for each locus averaged across 740 replicates and then all averaged together. Model parameters in Table S7C.

**Supplemental Figure 12.**
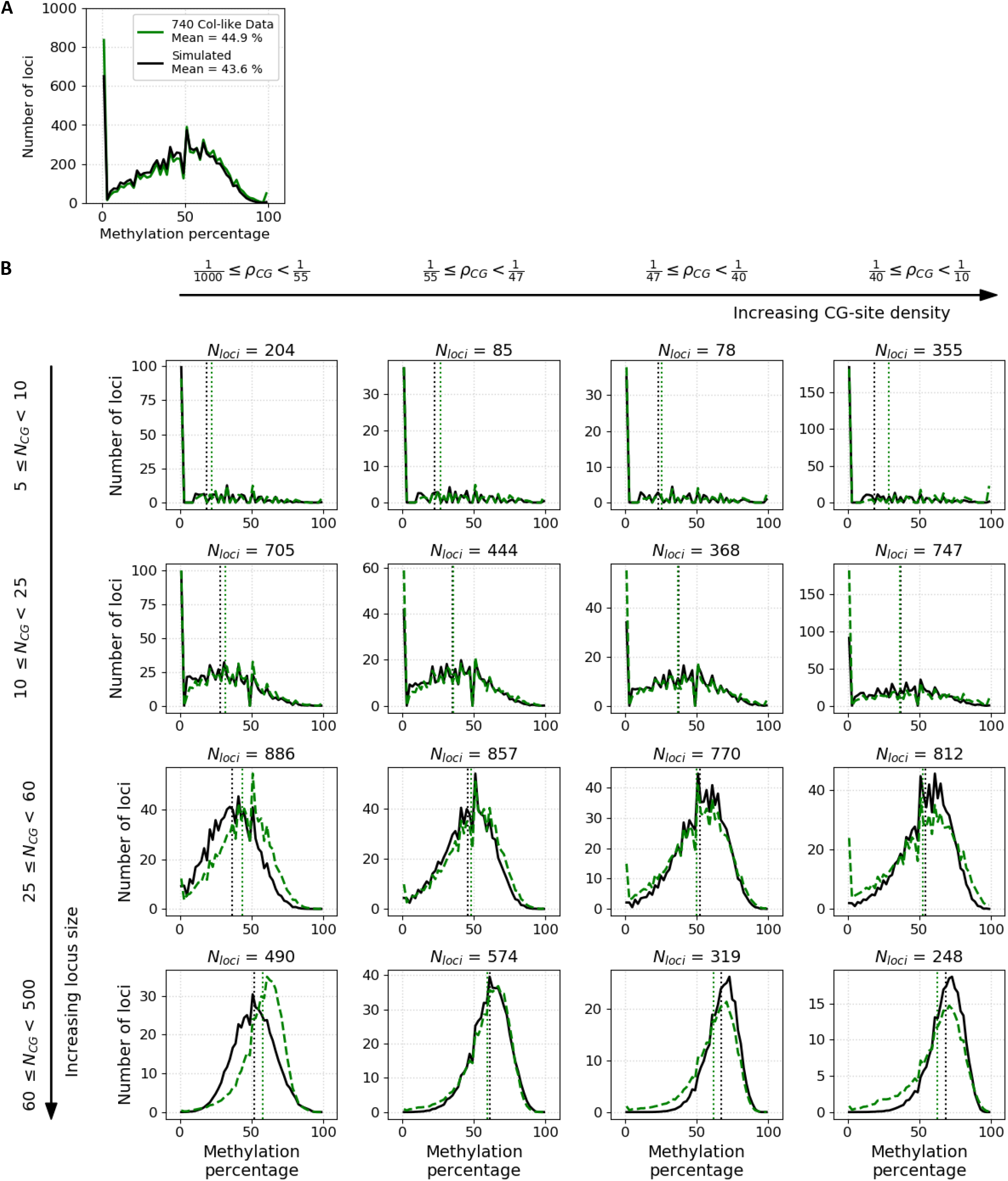
**A)** Distribution of methylation levels over methylatable regions for Col-like accessions (N=740, green) and from simulated steady-state (black). Simulations of full model run over 100,000 generations, normalized for 740 locus replicates, starting from all-*U* initial state. Model parameters in Table S7C. **B)** Distributions of methylation levels over methylatable regions for loci split by length (in CG sites, *N*_*CG*_) and average CG density (*ρ*_*CG*_) for Col-like accessions (N=740, green) and from simulated steady-state (black). Number of loci for each panel indicated and distribution means shown by dotted vertical lines. Simulations of full model run over 100,000 generations, normalized for 740 locus replicates, starting from all-*U* initial state. Model parameters in Table S7C.

**Supplemental Figure 13.**
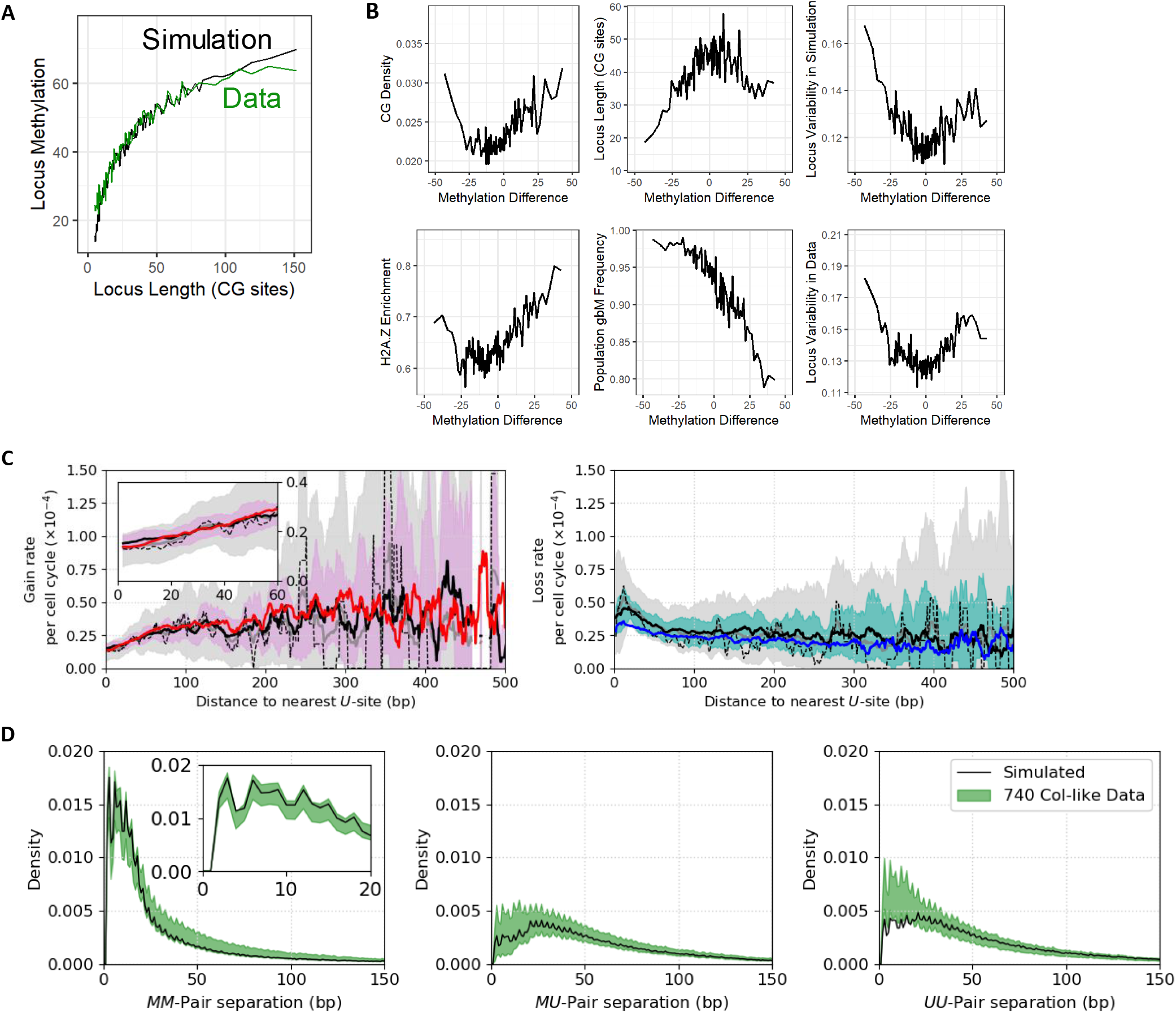
**A)** Longer loci are more highly methylated in both full model simulation results (black) and in the 740 Col-like accessions data (green). Loci were grouped into percentiles by CG site number, with 99% of percentiles shown and experimental methylation level averaged over N=740. Simulated methylation level for each locus was averaged over 740 replicates after 100,000 generations starting in all-*U* initial state. Model parameters in Table S7C. **B)** Full model performs less well for genes with high CG density, shorter length, higher H2A.Z level or those with low (or very high) population gbM frequency. Loci that are more variable (in the simulation or in experimental data) are also predicted less well. X-axis values calculated by subtracting the experimental methylation level of the locus (averaged over 740 Col-like accessions as in **A** from the simulated methylation level (simulated as in **A**), meaning that loci range from most under-methylated by the model to most over-methylated by the model. Loci were grouped into percentiles of methylation difference. **C)** Profiles of methylation gain (left) and loss (right) rates per cell cycle, plotted as a function of distance to the nearest uCG site, calculated over methylatable regions from full model. Experimental datasets labelled as described in Fig. 3E,F. Rates are shown as a 10 bp moving average. Inset highlights the gain rate at short length-scales. Model parameters, given in Table S7C, were not fit to these gain/loss rate profiles. Simulations performed as in Fig. S10A. **D)** Predicted (black) and observed (green) distributions of neighboring CG site separations are in good agreement (mCG-mCG (left), uCG-mCG (middle), uCG-uCG (right)). Full model simulation results normalized for 740 locus replicates after 100,000 generations starting from all-*U* initial state. Model parameters in Table S7C.

**Supplemental Figure 14.**
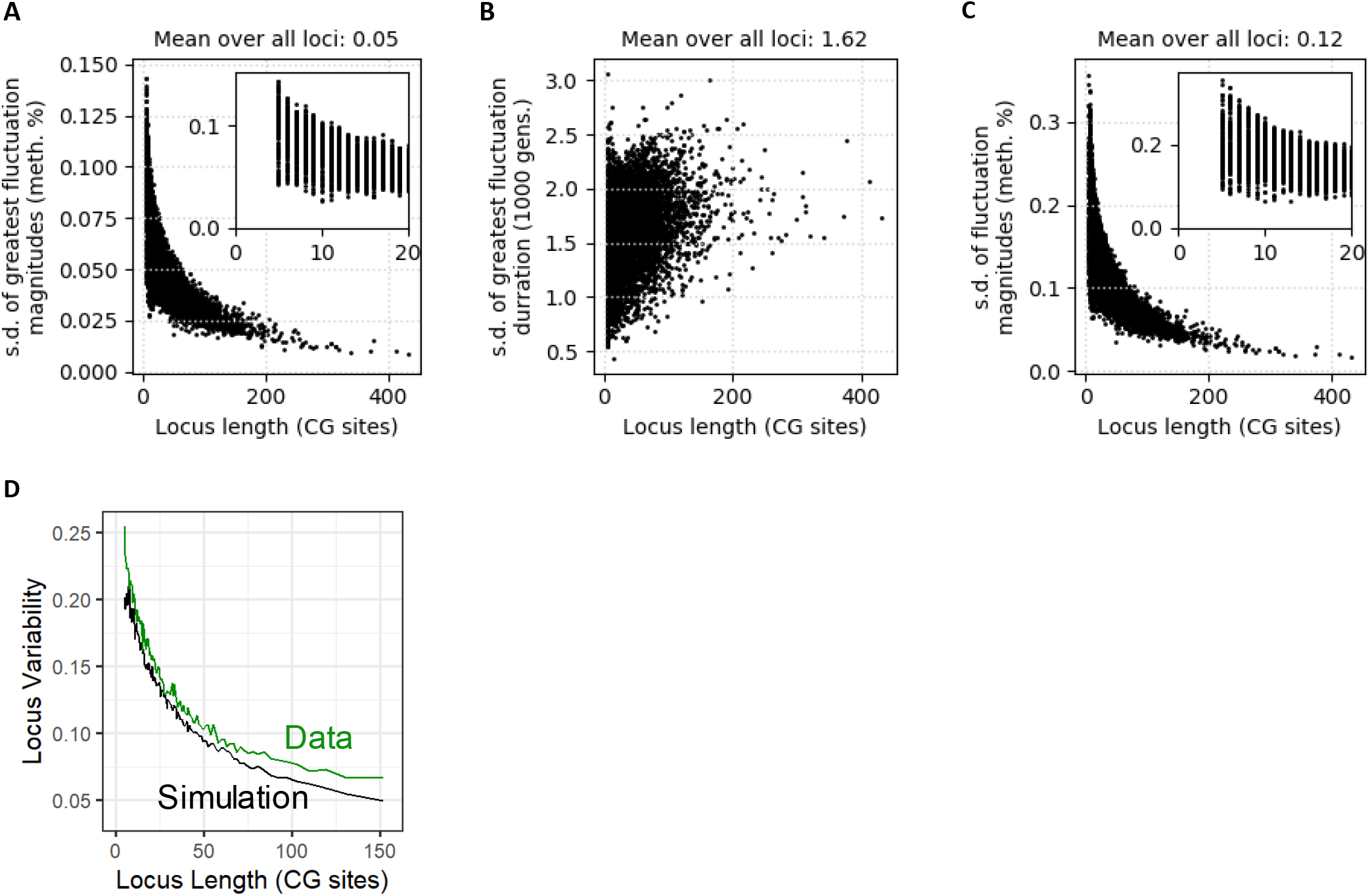
**A)** Standard deviation (s.d.) of magnitude of greatest overall locus methylation fluctuation from steady-state for loci of different lengths, using 740 locus replicates. Full model simulated for 100,000 generations starting from all-*U* initial state. First 50,000 generations used to equilibrate to steady-state. Second 50,000 generations used to measure fluctuations. Model parameters in Table S7C. **B)** Standard deviation of duration in generations of greatest magnitude methylation fluctuation from steady-state for loci of different lengths, using 740 replicates. Simulation performed as in **A**. Time shown in 1000 generations. **C)** Typical magnitude of methylation fluctuations away from steady-state (calculated by s.d. of fluctuations, Methods) plotted as a function of locus length. Simulated as in **A**. **D)** Shorter loci are more variable in both full model simulated results and accession data. Standard deviation of overall methylation levels of individual loci between 740 different Col-like accessions (Data, green) or different simulated realizations (Simulation, black) shown on y-axis. Loci were grouped into percentiles of length, with 99% of percentiles shown. Simulations run for 100,000 generations starting from all-*U* initial state with 740 realizations for each locus. Model parameters in Table S7C.

**Supplemental Figure 15.**
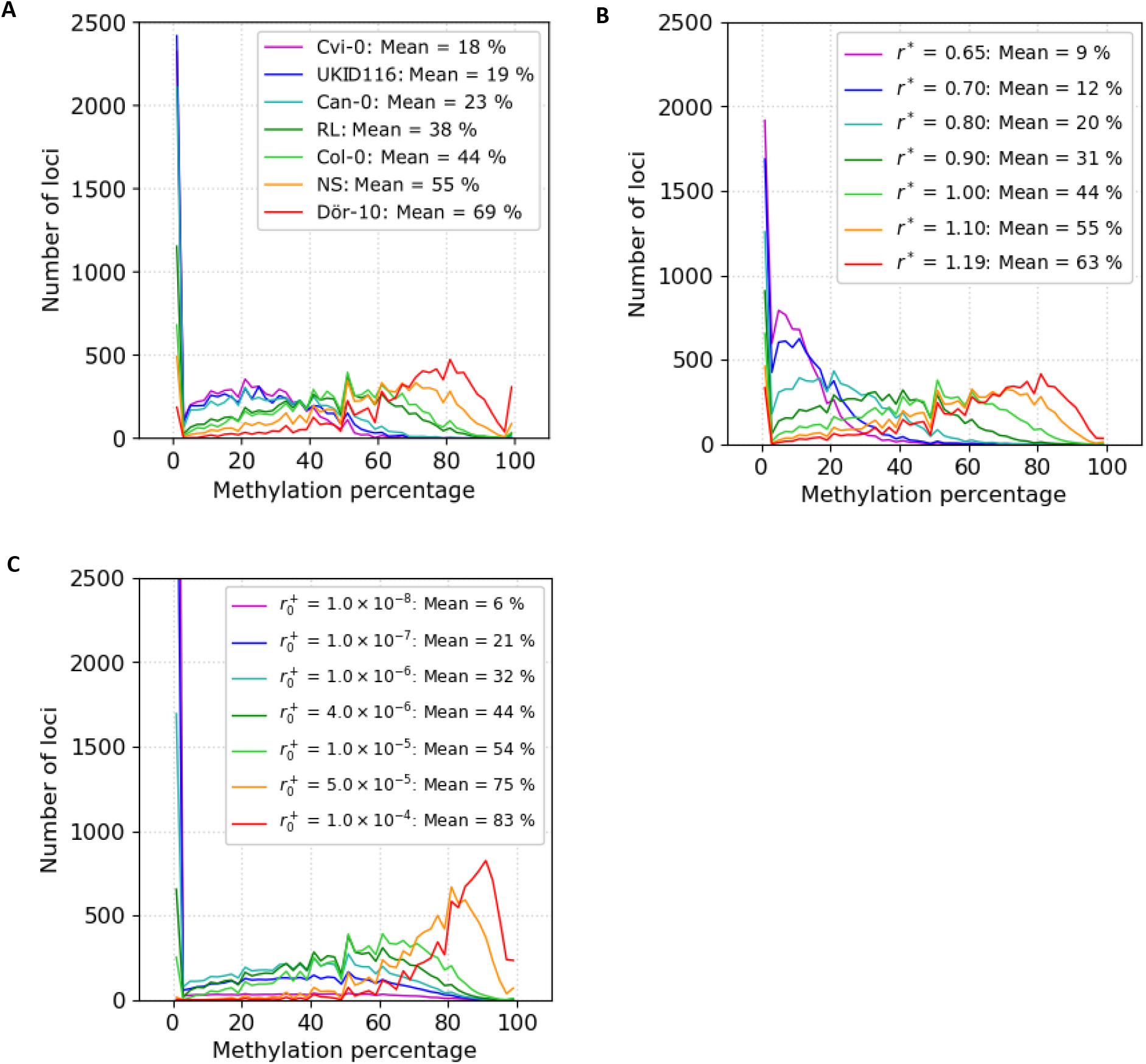
**A)** Distribution of methylation levels for loci within outlier accessions and Col-0. The mean for each dataset is included. **B)** Distributions of methylation levels produced by adjusting the strength of the cooperative interaction (*r*^∗^, Methods). The mean for each distribution is included. Simulated as in Fig. 6A. **C)** Distributions of methylation levels produced by adjusting the level of spontaneous *de novo* activity (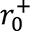, Methods). The mean for each distribution is included. Simulated as in Fig. 6A.

**Supplemental Figure 16.**
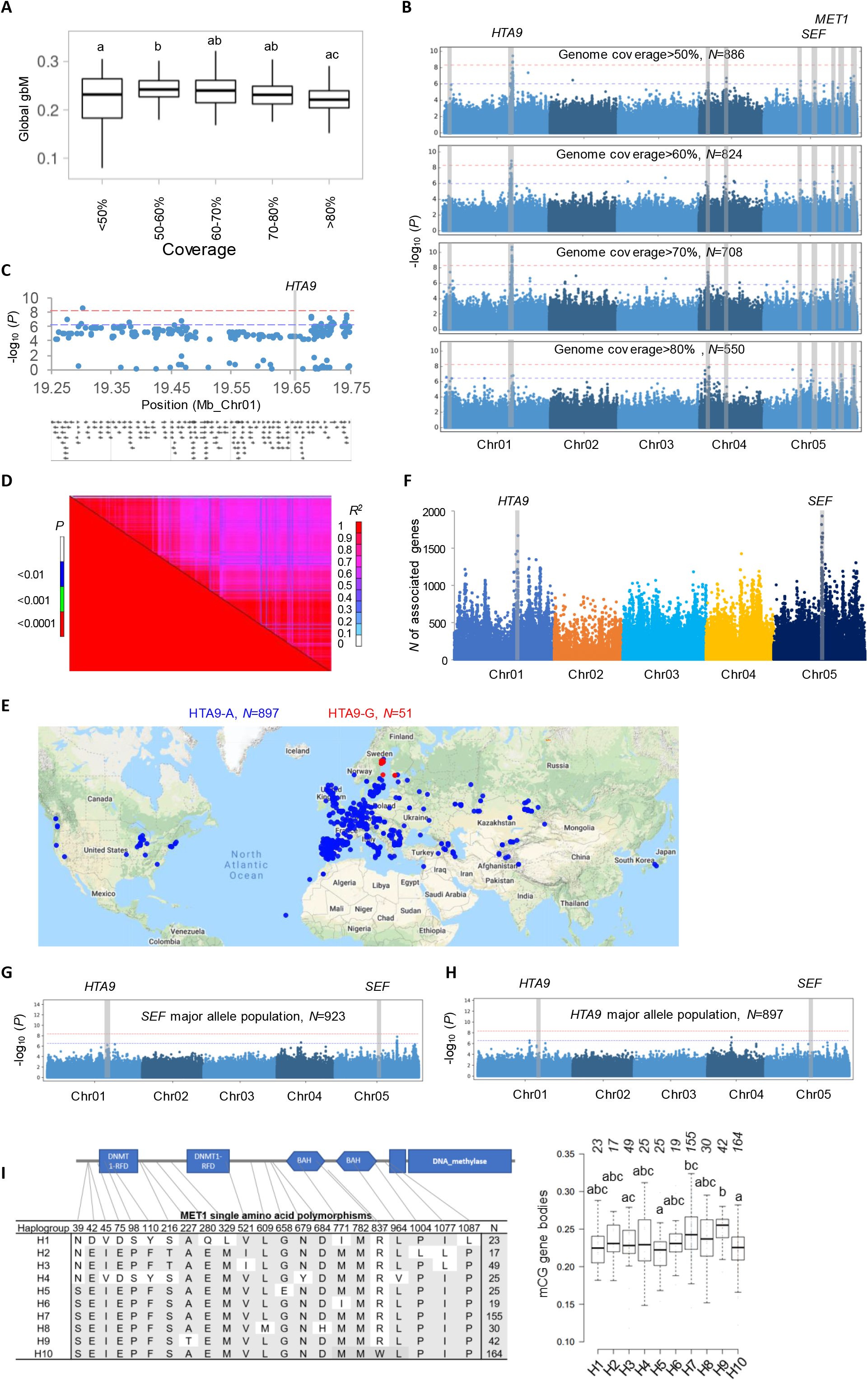
**A)** Global gbM levels are somewhat influenced by genome-wide sequencing coverage. Global gbM levels of accessions in different genome-wide sequencing coverage bins are plotted. Letters denote significant differences at p<0.05, one-way ANOVA, Tukey post hoc test. For example, groups denoted with “a” are statistically different from “b” and “ab” groups are different from neither “a” nor “b”. **B)** *HTA9, SEF* and *MET1* quantitative trait loci (QTLs) were detected at False Discovery Rate 0.05 (dashed blue line) at >50% genome coverage. *HTA9* was the only association detected at the Bonferroni threshold and robustly identified by GWA mapping using panels of accessions with >50%, >60%, >70%, and >80% genome coverage. Horizontal dashed blue and red lines respectively show the 5% False Discovery Rate and Bonferroni thresholds. **C)** The *HTA9* QTL is broad, covering a region of almost 500 kb. **D)** Linkage disequilibrium analysis revealed that SNPs in this 500 kb region covering HTA9 association display long range linkage disequilibrium. **E)** Geographic distribution of *Arabidopsis* accessions, with accessions containing the *HTA9-A* allele shown in blue and those containing the minor *HTA9* allele (*HTA9-G*) in red. **F)** *HTA9* and *SEF* were again identified as significant QTLs from GWA mapping using intragenic CG methylation levels of individual genes in the 1001 methylomes data. Multiple linked SNPs in HTA9 genomic region were associated with gbM variation of a total of 4074 genes at P<0.0001 (accelerated mixed model). Associated SNPs in HTA9 and SEF regions are in linkage disequilibrium (R^2^ =0.21, standardized disequilibrium coefficient (D’) =0.66, P=1.2 x 10^-17^, two-sided Fisher’s exact test). **G)** *HTA9* still displays strong association (P=5.12 x 10^-7^, mixed model regression) with gbM after accounting for the SEF genetic polymorphism. Horizontal dashed blue and red lines respectively show the 5% False Discovery Rate and Bonferroni thresholds. **H)** *SEF* no longer displays strong association with gbM after accounting for the *HTA9* genetic polymorphism, indicating that the SEF association with gbM is likely spurious and detected due to linkage disequilibrium with HTA9 SNPs. Horizontal dashed blue and red lines respectively show the 5% False Discovery Rate and Bonferroni thresholds. **I)** Position and nature of amino acid polymorphisms among *MET1* haplogroups. The accessions having identical amino acid sequences are classified into a haplogroup, and haplogroups with a minimum of 15 accessions were retained for this analysis. The number of accessions within each haplogroup are indicated. The right panel shows boxplots for gbM of accessions belonging to each MET1 haplogroup. Letters denote significant differences at p<0.05, one-way ANOVA, Tukey post hoc test.

